# Ancient Mitogenomes from Belize Provide Insights into the Mobility and Migration of Ancient Communities in Mesoamerica and the Caribbean

**DOI:** 10.64898/2026.10.06.756460

**Authors:** Celia D. Cleary, Lauren C. Springs, Angelina J. Locker, Diane Z. Chase, Arlen F. Chase, Adrian S. Z. Chase, Melissa M. Badillo, Adela Pederson Vallejos, Genara Cano, Roy Rodriguez, Humberto García-Ortiz, Lorena Orozco, Jada Benn Torres, Rick W.A. Smith, Austin W. Reynolds

## Abstract

Maya peoples have undergone several largescale demographic transformations over the past 3000 years, including the formation and depopulation of large urban centers, climatic events, and multiple colonial regimes. Archaeological and stable isotope evidence from across the Maya lowlands throughout these periods paints a picture of a demographically mobile population with complex and dynamic relationships among some of the largest urban polities in antiquity. The nature of the genomic relationship between ancient Maya urban centers, as well as their relationship with present-day Maya communities, however, remains under-characterized. Here, we present mitogenomic data of 16 ancient Maya Ancestors recovered from interments in the ancient Maya cities of Caracol and Santa Rita Corozal located in Western and Northern Belize. These ancient data were combined with 110 newly collected contemporary individuals from Northern Belize, along with nearly 600 previously published ancient and contemporary comparative mitogenomes recovered from sites spanning Mesoamerica and the Caribbean to reconstruct paleodemographic and mobility patterns of maternal lineages through time. Our results are consistent with broadly interconnected Maya populations that have maintained maternal genetic continuity across much of the Holocene, and which are not structured by linguistic family.

**Significance:** Recent developments in genome-wide studies have provided valuable insights into the relationships between Indigenous Peoples across Mesoamerica. Fewer have focused on matrilineal relationships specific to the Maya, a group that has had a sustained influence in the region for millennia and had some of the largest urban polities since antiquity. Here, we contribute the whole mitogenome sequencing of 16 ancient Maya Ancestors from the polities of Caracol and Santa Rita Corozal in Belize. We additionally contribute 110 whole mitogenomes from living residents who identify as Maya people living in the Northern Belizean district of Corozal, a region known to have a complex migratory history, past and present. We find evidence of sustained maternal lineage sharing both between Maya-speaking groups, and with other groups across Mexico, a pattern not clearly structured by linguistic affiliation. This study contributes to characterization of the complex demographic history in Mesoamerica by providing diachronic population genomic analysis at a more granular level.

## Introduction

Genomic analyses of Indigenous and Indigenous-descended peoples have revealed complex patterns of genetic variation shaped by similarly complicated demographic histories, geography, ecology, and colonial events^1–8^. Understanding the regional-scale relationships among these peoples is important for accurately interpreting these genomic patterns and reconstructing the demographic processes that produced them. This is especially important for Indigenous peoples of the Americas, whose genetic diversity is not only shaped by the complexities of its initial peopling and continuous demographic changes, but also constitutes a mosaic of European, African, and Asian ancestries introduced since the 16^th^ century^1,3,8,9^. Recent genomic studies show that these admixture histories vary widely geographically and temporally ^2–4,6,10,11^. Other studies indicate the importance of the relationship between ancient population history to present-day genetic variation, showing both patterning consistent with regional ecological zones and recapitulation of population structure established before European colonialism among the Indigenous ancestry tracts of Mestizo peoples ^2,3^.

Among Central American populations, populations that fall within the Maya linguistic family have undergone particularly significant demographic transformations from ancient to present-day, with demographic transformations including the formation and depopulation of major urban centers, climatic events, and extensive networks of trade and exchange in addition to multiple regimes of European colonialism ^12–24^. Though radiocarbon dating shows that people occupied the land known today as the Maya Lowlands at least since 9.5kya^24^, the first settlements definitively identified as Maya via broadly shared ceramic culture were in the Middle Preclassic (1000-400 BCE)^16,25–27^. These ceramic-producing cultures spread quickly throughout the Maya Lowlands, creating sedentary communities and elaborating the processes of social stratification^28,29^. The Classic Period (250-∼900 CE) saw major demographic growth paired with the development of some large urban centers which had extremely dynamic relationships with one another in terms of allyship, war, and even imperialism, resulting in similarly dynamic migratory patterns across the region and a high degree of gene flow^15–17,24,30–32^.

The demographic and political decline that followed during the Terminal Classic Period (∼750 CE-900 CE) had extremely varied effects across Maya sites and their peripheries with some polities experiencing depopulation while others grew^33,34^. These effects further altered migratory patterns as people moved away from areas of political instability and war to others which were more buffered from the effects of demographic and political change^12,13,16,35–37^. British and Spanish colonialism also resulted in major demographic transformation among Maya groups and was also heterogeneous in its effects^38–40^. Although a myriad of archaeological and historical data provide evidence for high levels of migration in this region throughout all these periods, much less is known about the impact of these events on the genetic relationships between Maya groups over time.

Recent genetic studies have begun to document the genetic history of Maya populations^1,11,24,41–44^, though the nature of the genomic relationship between ancient Maya urban centers, as well as their relationship with present-day Maya communities, remains under-characterized. Genome-wide studies place this history in the context of the peopling of the American double-continent, providing a picture of Mesoamerican demographic history as a rapid dispersal across the region with subsequent differentiation of groups that maintained some degree of gene flow between one another over time^1,11,43^. Such studies have modeled genetic divergence between Northen and Central Mexican populations, such as the Tarahumara and the Huichol, from Mesoamerican populations, such as the Triqui and Maya, occurring towards the beginning of the Holocene (∼11.7kya-present)^1,43,45^. The most diverged groups in the Mesoamerican clade, the Triqui and the Maya, are estimated to have split around 5.5kya^1^, followed by a South-to-North migration from genetically distinct Isthmian populations into Belize in a between 5.5 and 3kya ^24^.

Early studies of mtDNA and NRY haplogroups, along with autosomal STRs provide limited information regarding the relationship between Maya populations, suggesting that groups sharing the Maya linguistic family (composed of about 31 different languages) share a common genetic history and are distinct from other Indigenous groups in Mesoamerica and South America^43,46–48^. Although closely related to one another, these haplogroup-level studies indicate a high degree of genetic heterogeneity between Maya populations indicative of sub-structuring in the Maya clade^42,47–49^. The addition of ancient DNA (aDNA) into studies of Maya population history, which until now have largely focused on population size and growth^50^, have begun to unravel the relationship between ancient and present-day Maya populations. These studies note a fair amount of genetic continuity in certain polities from the Classic Period to present-day Maya towns in the local vicinity^51^, as well as far reaching genetic connections between Classic Period cities that went beyond the Maya region^44^.

Demographic transformations occurring during colonialism further complicate the genetic relationships between ancient and present-day Maya populations, although genome-wide investigation into these colonial processes have begun to provide additional insight into these dynamics. Many Maya people today have large portions of European and African genetic ancestry, due to Spanish and English colonization and the Transatlantic Slave Trade. Gene flow during the Colonial Period was heavily sex biased, with more Indigenous and African genetic ancestry being passed down through maternal uniparental markers (mtDNA) and European ancestry being passed through paternal uniparental markers (Y chromosome)^3^. Fragment length distributions of African genetic ancestry in living Central Americans indicates two major pulses of migration during the Early Colonial Period (one pulse ∼14 generations ago and another ∼12-5 generations ago), consistent with the timing of the Transatlantic Slave Trade^3^. In addition, Maya revolt against Spanish colonial rule in 1847, known as the Caste War, saw nearly half the population of the Yucatán flee the area in the years following, resulting in the movement of many Maya people south into Northern Belize^52^. Together, these findings suggest that Maya population history is complex, and there is still much to learn about the genomic relationships between living Maya peoples, the relationships between the ancient and contemporary Maya, and the genomic consequences of colonialism in the Maya area.

In this study, we combine previously published mitochondrial genomes from ancient and living individuals with newly collected ancient and contemporary mitogenomes to understand how present-day Maya peoples are related to one another, how present-day Maya peoples are related to ancient Maya peoples, and to investigate recent admixture in the maternal lineages from outside of the Americas. Our results indicate broad maternal lineage sharing with increasing genetic differentiation between ancient and present-day Maya peoples as time-depth increases, consistent with maternal lineage sharing between Maya groups extending since at least the Classic Period (250-∼900 CE)^16^.

## Results

This study generated 16 novel whole mitogenome samples from the ancient Maya urban centers of Caracol and Santa Rita Corozal, along with 110 whole mitogenomes from living Maya people from the Northern Belize district of Corozal (**Table S1**). Ancient mitogenomes generated here were recovered from residents of two Maya cities: Caracol and Santa Rita Corozal. Caracol was one of the largest urban centers in antiquity, reaching its peak occupation in the Classic Period (250 CE – 900 CE); it was unoccupied after the Classic Maya collapse (by 910 CE) (**Table S1**)^53^. Individuals recovered from Santa Rita Corozal exhibited A1 (11.11%), A2 (66.67%), B2 (11.11%), and C1 (11.11%) maternal haplogroups, with A2 being the most common at the site. Caracol individuals exhibited only A and A2 haplogroups, again with A2 as the most common haplogroup **(Fig. 1A)**.

**Figure 1.**
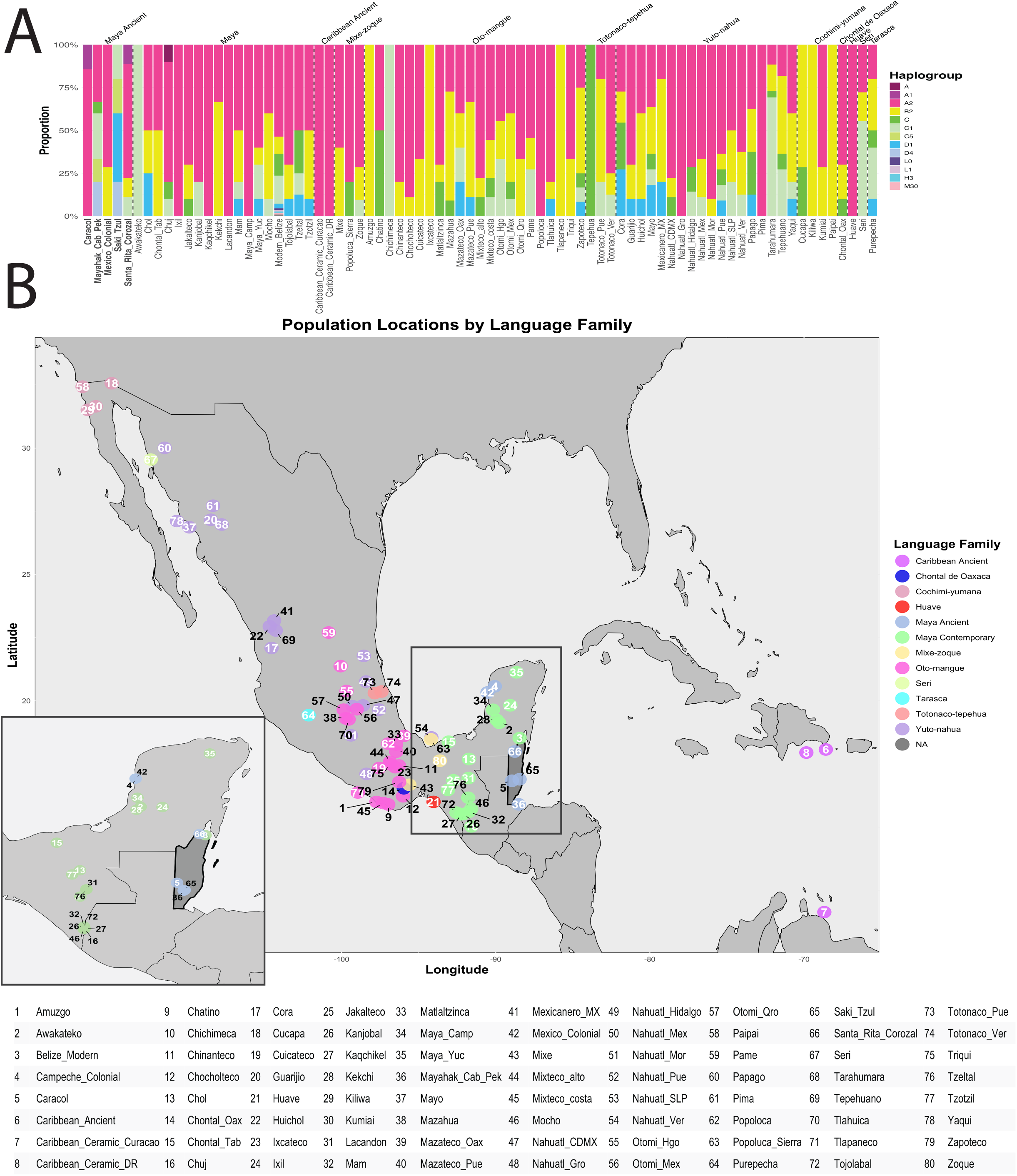
A) Haplogroup distribution per population grouped by language family; B) Map showing samples used in this study. Numbers indicate population, while colors indicate language family of the individuals or their ancestors. Pop-out map shows Maya speakers included in this study.

Present-day whole mitogenomes generated by this study came from living residents who identify as Maya in 9 villages in the northern Belizean district of Corozal (**Fig 1**, **Table 1, Table S1)**. These participants exhibited A2, B2, C, C1, and D1 Indigenous American haplogroups, as well as four maternal haplogroups originating from outside the Americas: L0, L1, H3, and M30. These data were combined with previously sequenced present-day mitogenomes from the MAIS cohort^11,54–56^ along with Archaic Period (8000 – 2000 BCE) mitogenomes from Mayahak Cab Pek and Saki Tzul, and Colonial Period (beginning 1500)^16^ mitogenomes from the Spanish-established colonial town of Campeche, resulting in a total sample size of 724 (**Fig 1**, **Table 1, Table S1**).

Initial assessment of haplogroup composition of the entire dataset shows that maternal haplogroup A2 (53.6%) is by far the most common haplogroup across our study area, consistent with broader patterns across North and South America. Notable exceptions to this trend are the Archaic-Period individuals from Mayahak Cab Pek and Saki Tzul, as well as contemporary Seri, Cora, Tarahumara, Tepehua, and Purepecha groups, where haplogroups C, C1, C5, D1, and D4 were more common (**Fig. 1**).

### Haplogroup Composition Varies Through Time in the Maya Region

Wrights’ fixation indices (F_ST_) of Indigenous haplogroups representing pairwise genetic differentiation between Mayan-speaking populations show little maternal lineage differentiation between groups broadly, with the exception of the most ancient groups (Caracol, Mayahak Cab Pek, Saki Tzul) and some present-day Maya groups previously reported to have high levels of genetic drift (Kaqchikel, Lacandon)^43^. Comparisons of living Maya people in Corozal District, Belize and other living Maya groups (Kekchi, Campeche, and Yucatan) show little to no differentiation (F_ST_ = 0.113, p = 0.086±0.003; F_ST_ = 0.027, p = 0.246±0.005; F_ST_ = 0.000 p = 0.364±0.005, respectively) **(Fig. S2)**. Multi-Dimensional Scaling (MDS) based on pairwise F_ST_ values show no distinct clusters either in the dataset as a whole (**Fig.2A**) or in a subset of data including only those individuals in the Mayan language family (**Fig. 2B**). Fisher’s exact tests measuring differences in haplogroup proportions between each pair of populations also show little maternal differentiation between Maya groups. The only present-day Maya groups observed to have small but significant haplogroup differentiation from the present-day Belizean Maya population from Corozal were Tzotzil (p=0.032) and Mocho (p=0.045), who were both sampled in the Mexican state of Chiapas.

**Figure 2.**
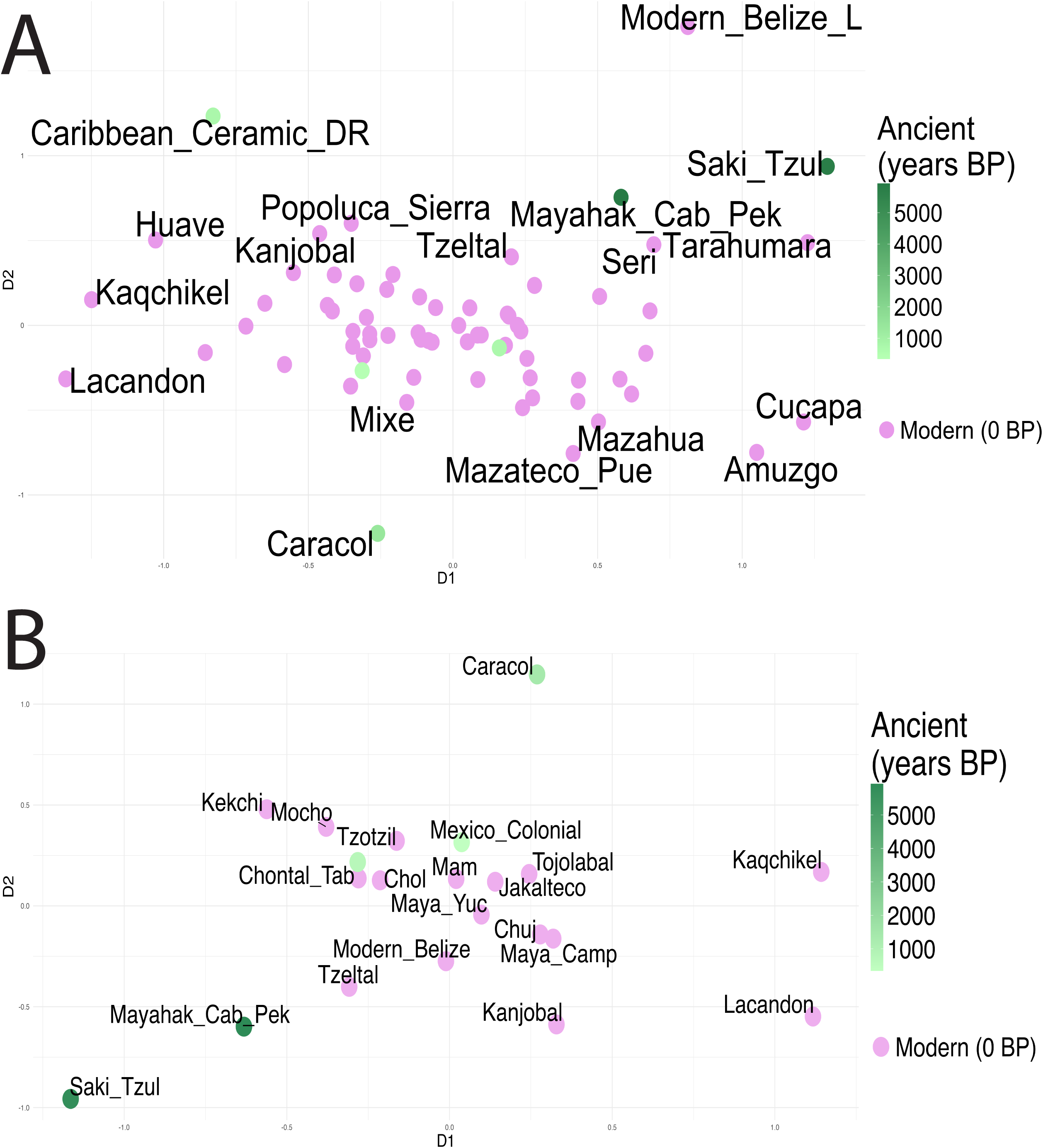
MDS showing A) all study populations, and B) Maya speakers.

Chi-squared (χ^2^) tests were used to estimate over- or under-representation of specific haplogroups in each group, as compared to the entire comparative dataset. Haplogroup C exhibited over-representation in three of the Indigenous Mexican groups that were not in the Maya language family, namely C1 in Seri (C1 = 5.122, p = 2.455*10^-5^) and Tarahumara (C1 = 8.118, p = 8.478*10^-14^), located in the Mexican state of Chihuahua, and C (nonspecific) in the Tepehua (C = 3.774, p = 0.010), located in Jalisco. The present-day Chuj also exhibited an over-representation of haplogroup A (nonspecific) (A = 8.415, p = 8.745*10^-15^). The present-day Belizean Maya from Corozal show no significant over- or under-representation of any of the maternal haplogroups observed across the entire dataset (**Fig. S4; Table S9**).

A median-joining network of maternal haplogroup A shows the least amount of haplogroup diversity despite having the highest representation in the dataset and displaying the expected “star-shaped” phylogeny characteristic of a founder event followed by rapid expansion of a lineage^57,58^. Present-day Maya people tend to be placed in the center nodes of the network with relatively large portions of the individuals included in the network sharing the same few haplogroups: A2+(64) (11.06%), A2 (8.54%), A2m (6.03%), and A2w (6.03%) (**Fig. 3**). Networks for maternal haplogroup B again show the expected star-shaped phylogeny observed previously^57,58^. Present-day Maya mitogenomes tend to pair closely with Yuto-nahua and Oto-mangue speakers, with the present-day Maya from Corozal being spread throughout the network. The most common haplogroups in this network are B2 (19.35%), B2t (9.66%), and B2+16278 (6.21%) (**Fig. 3)**. Most common maternal haplogroups in the networks for haplogroup C were nonspecific C (28.28%), C1c1a (14.14%), C1d1 (14.14%), C1c5 (12.12%), C1c (10.10%), and C1c4 (5.05%) (**Fig. 4**). The network for maternal haplogroup D shows one central node from which all others radiate. Present-day Maya mitogenomes in the maternal haplogroup D network mostly radiate from the ancestral Indigenous D haplotype, sometimes pairing with Oto-mangue-speakers. The most common maternal haplogroups in this network are D1 (40.63%), D1f3 (9.38%), D1i2 (9.38%), D1m (9.38%), D1h1 (6.25%), D4h3a (6.25%), and D4h3a5 (6.25%) (**Fig. 4**).

**Figure 3.**
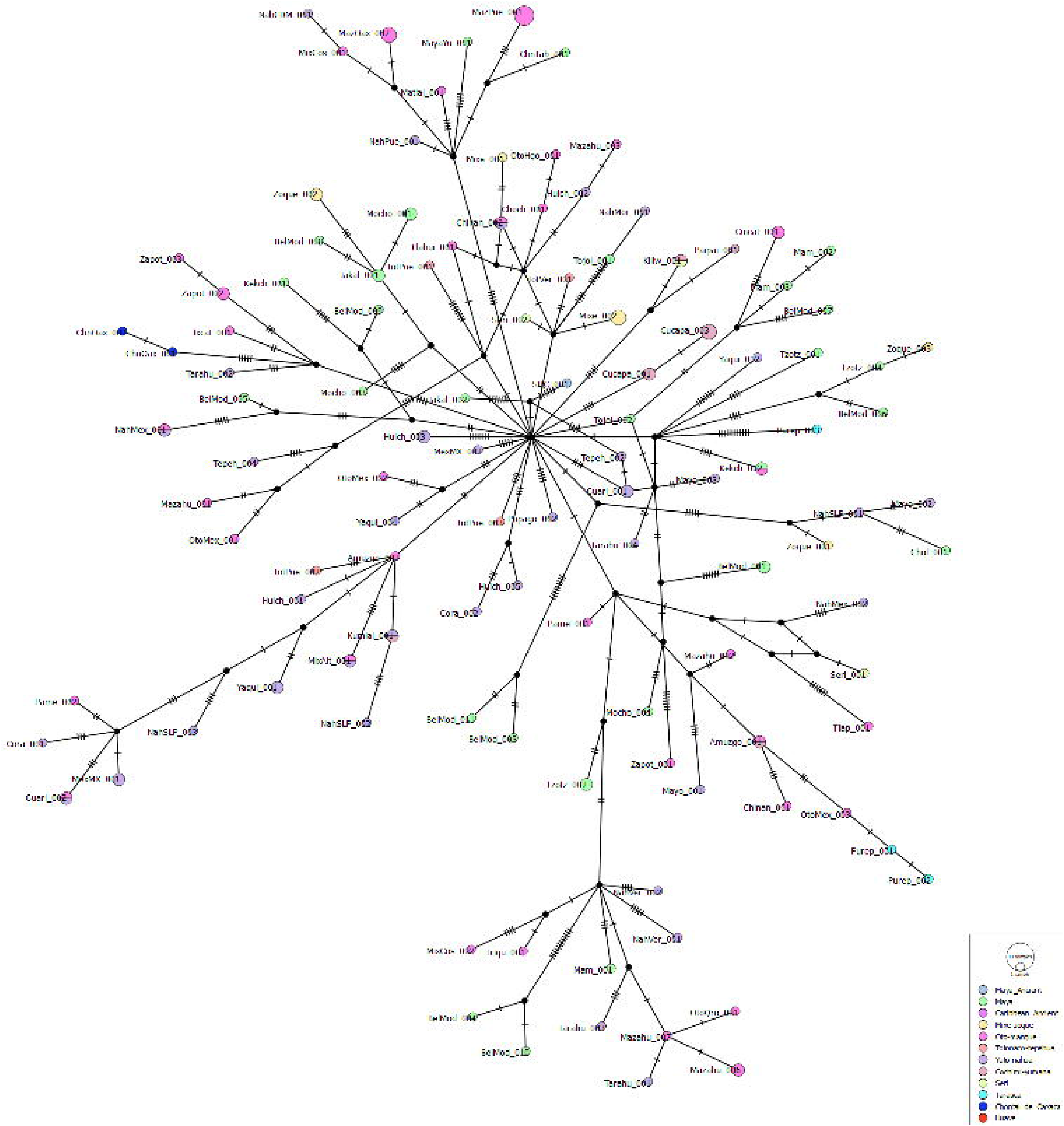
Haplogroup A median joining network. Size of nodes indicate number of samples with that maternal haplogroup, while color indicates population or proportion of samples represented by each population. Tick marks are proportional to the number of molecular differences between haplotypes. Loops represent alternative, equally parsimonious connections between haplotypes.

**Figure 4.**
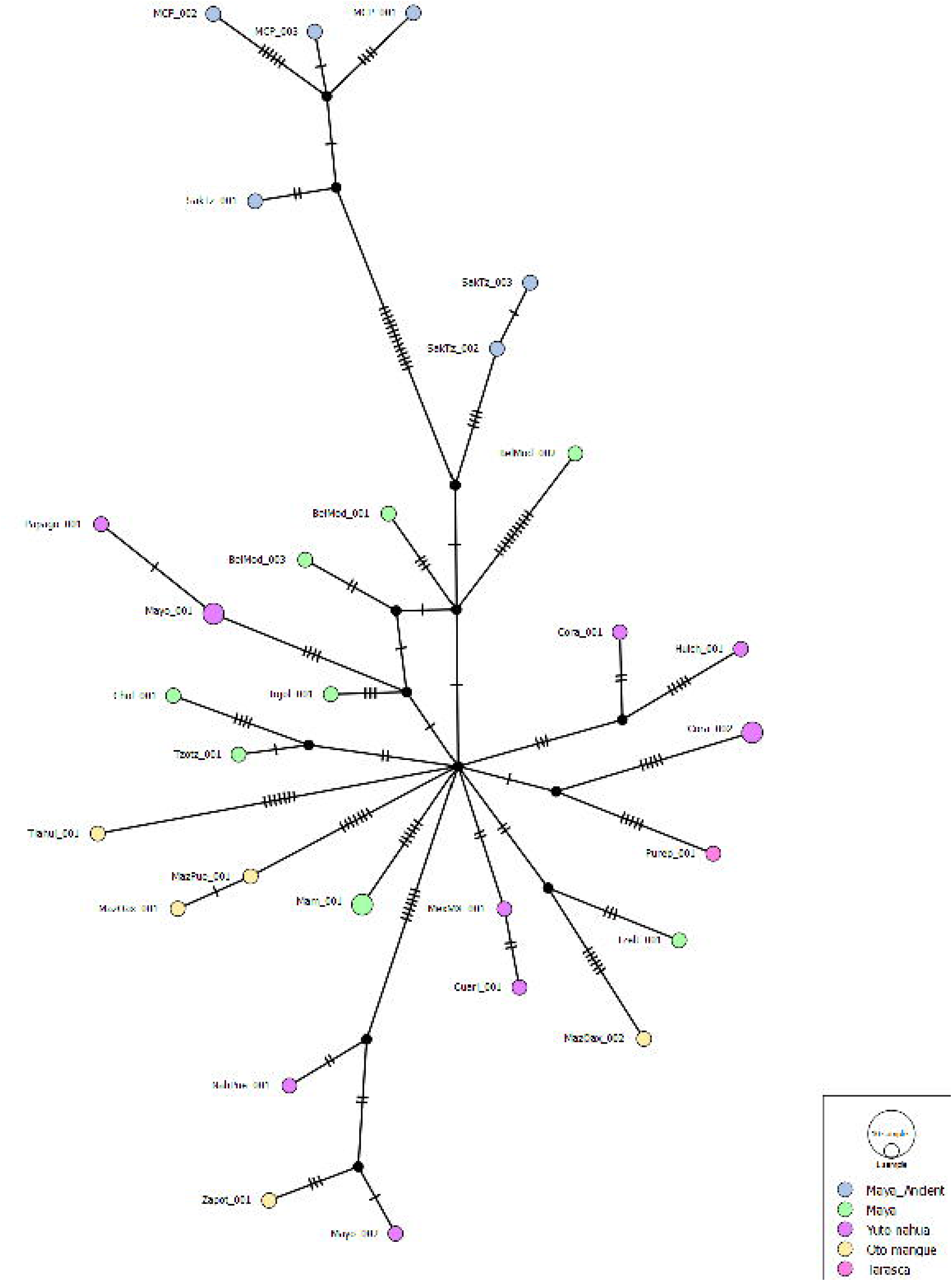
Haplogroup C median joining network. Size of nodes indicate number of samples with that maternal haplogroup, while color indicates population or proportion of samples represented by each population. Tick marks are proportional to the number of molecular differences between haplotypes. Loops represent alternative, equally parsimonious connections between haplotypes.

Bayesian phylogenetic analysis of all ancient and present-day individuals in this study with the maternal haplogroup A showed that while many living Maya people (Mayan-speakers and those living today who are descended from Mayan-speakers) clustered together, others were disbursed throughout the tree. Haplogroup clades in which multiple language groups are represented show some additional affinity of Mayan-speakers towards Yuto-Nahua, Oto-Mangue, and Mixe-Zoque speakers (**Fig. 5A**). Bayesian phylogenetic analysis of individuals with maternal haplogroup B showed a similar trend, with present-day Maya speakers appearing in nearly every major clade. Mayan speakers and their descendants in mixed-language clades showed a tendency to cluster with Yuto-Nahua, Oto-Mangue, and Mixe-Zoque speakers, possibly reflective of geographic proximity (**Fig. 5B**). This finding is corroborated by a Mantel test, which revealed a significant positive correlation between pairwise F_ST_ and geographic distance (r = 0.255, p = 0.005, 9999 permutations) consistent with higher levels of isolation by distance among sampled groups that are closer to each other geographically **(Fig. S7)**. Present-day Maya individuals living in Corozal appeared to cluster together as well, possibly reflective of geographic proximity or of common migration from the Yucatán **(Fig. 5B)**. Bayesian phylogenetic analysis including all individuals in the dataset with maternal haplogroup C showed few individuals from the Mayan language family. No present-day Belizean Maya individuals appeared in this tree (**Fig. 5C**). Bayesian phylogenetic trees for all individuals with maternal haplogroup D showed present-day individuals in the Maya language family clustering together in a clade with many of the Yuto-Nahua speakers. Only two present-day Maya individuals from Corozal appeared in this haplogroup (**Fig. 5D**).

**Figure 5.**
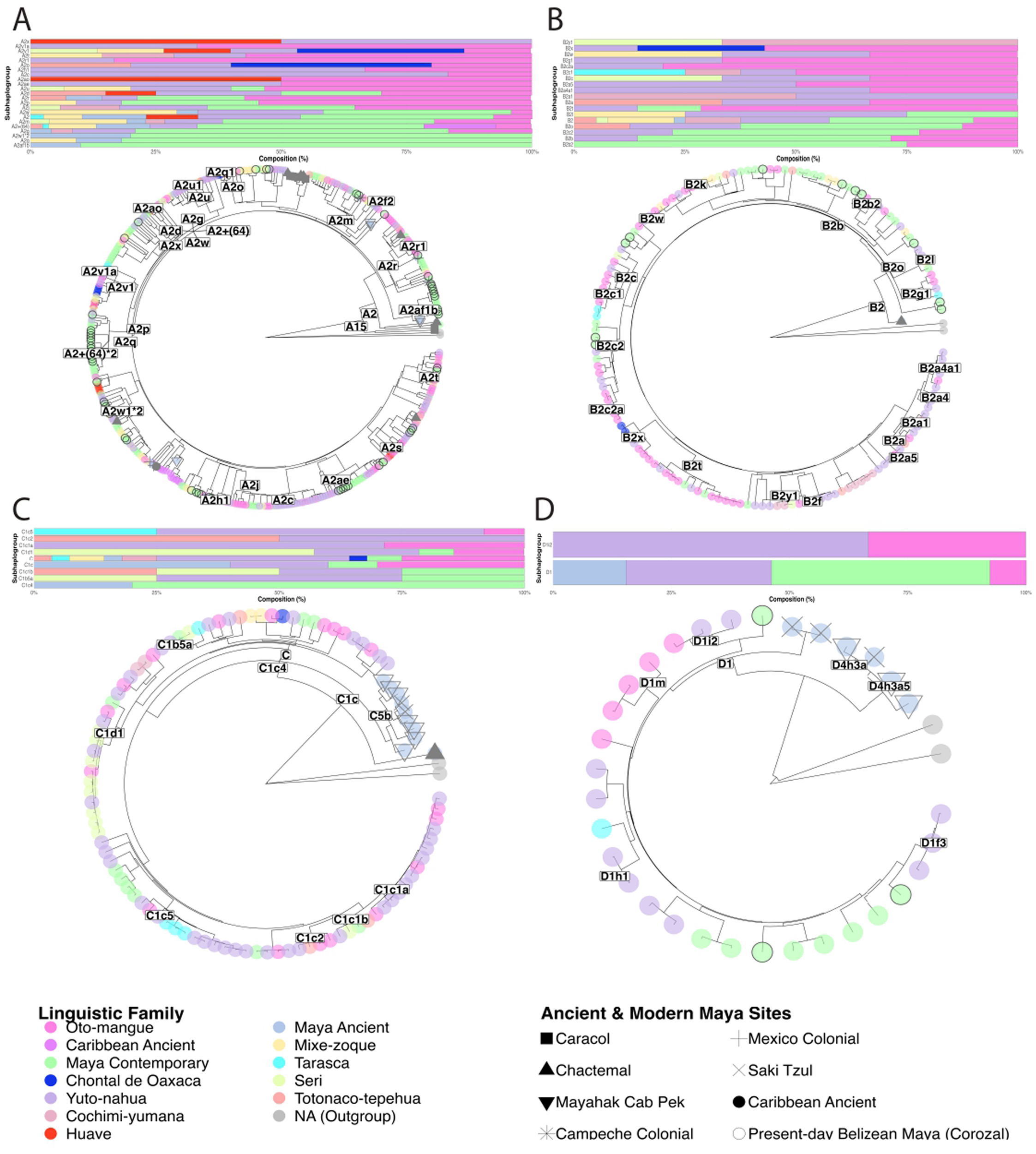
Bayesian phylogenetic trees for A) Haplogroup A, B) Haplogroup B, C) Haplogroup C, and D) Haplogroup D. nodes are colored based on language family. Present-day Belizean Maya from Corozal are denoted with black circular outlines, and ancient sites with shapes. Trees are accompanied by par plots showing the language group composition of clades containing more than one language group.

### Temporal Comparisons Reveal Changes in Haplogroup Composition over Time in the Maya Region

Comparisons of present-day Maya from Corozal and ancient Maya groups show a general trend of increasing differentiation with increasing mean temporal distance between groups. The most ancient individuals included in this study were pre-Maya, recovered from Saki Tzul (mean age = 5,768.4 BP; age range = 7,376 – 4,482 BP) and Mayahak Cab Pek (mean age = 5,925.667 BP; age range = 9,534 – 3,860 BP) rock shelters ^59,60^. For the sake of this study, those recovered from both of these rock shelters were grouped together with ancient Maya groups, due to their location within the Maya region. Not only was Saki Tzul significantly differentiated from present day Maya individuals in Corozal (F_ST_ = 0.246, p = 0.003±0.001), but the mitogenomes from this ancient site was also significantly differentiated from all other Maya groups except for Mayahak Cab Pek (F_ST_ = 0.036, p = 0.243±0.003) (**Fig. S1**). Mayahak Cab Pek was also found to be significantly differentiated from present-day Maya living in Corozal (F_ST_ = 0.104, p = 0.006±0.001). Caracol (mean age = 1,375 BP) was also differentiated from present-day Corozal Maya (F_ST_ = 0.111, p = 0.032±0.002) as well as nearly all other ancient and present-day Maya groups. The exception to this trend was found in Santa Rita Corozal. Though F_ST_ between Santa Rita Corozal and Caracol indicates moderate levels of differentiation (F_ST_ = 0.114), the comparison did not reach statistical significance. Santa Rita Corozal (mean age = 709 BP) showed virtually no differentiation from present-day Maya in Corozal (F_ST_ = 0.000, p = 0.9999±0.000). Interestingly, Santa Rita Corozal was significantly differentiated from Saki Tzul, but not Mayahak Cab Pek. Colonial Period samples from Campeche were not shown to be differentiated from present-day Maya in Corozal (mean age = 348 BP), with an F_ST_ of 0.078 (p = 0.059±0.002) (**Fig. S1**).

Though MDS shows no distinct clusters, there is some separation based on temporal distance. When all populations were included, Dimension 1 (D1) separated from the central point cloud Saki Tzul, which had the oldest mean age (mean age = 5,768.4 BP), as well as some of the present-day populations which have been affected by high levels of genetic drift (Kaqchikel, Lacandon, Huave, Tarahumara, Amuzgo, and Cucapa). Dimension 2 (D2) also drew ancient groups (Saki Tzul, Mayahak Cab Pek, Caracol, Caribbean Ceramic) out from the central cluster. When the MDS was limited to Maya-speaking populations only, D1 and D2 similarly isolated the oldest populations (Saki Tzul, Mayahak Cab Pek, Caracol) from the central cluster of populations, along with the present-day populations most affected by genetic drift (Kaqchikel, Lacandon) (**Fig 2**).

Fisher’s exact tests corroborate the finding that the oldest mitogenomes were the most differentiated from present-day Maya and Indigenous Mexican groups. Notably, out of the ancient Maya and the Corozal Maya mitogenomes generated in this study, the only group that is consistently differentiated from present-day Yuto-nahua-speaking populations or present-day Oto-mangue-speaking populations is ancient Saki Tzul. Similarly, Mayahak Cab Pek and Saki Tzul are the only groups consistently found to be separate from present-day Mixe-zoque speakers. Saki Tzul was observed to be differentiated at α=0.05 from most of the present-day Maya populations except Tzeltal (p=0.210), Maya from Yucatan (p=0.058), present-day Maya from Campeche (p=0.238), Chontal from Tabasco (p=0.522), and Chol (p=0.521). Mayahak Cab Pek was differentiated from all the other Maya groups except Tzeltal (p=0.356), Maya from Yucatan (p=0.253), present-day Maya from Campeche (p=0.795), Mam (p=0.063), Kekchi (p= 0.091), Kanjobal (p=0.169), Chuj (p=0.209), Chontal from Tabasco (p=0.350), and Chol (p=0.176). Early Colonial Period individuals from Campeche were not shown to be differentiated from any of the present-day Maya populations, and the only ancient Maya group differentiated from this population was Saki Tzul (p=0.007). The only Maya group found to be separate from Caracol based on haplogroup frequencies are Mocho (p=0.034) and Saki Tzul (p=0.004). Mayahak Cab Pek was not differentiated from any of the other ancient Maya groups. Santa Rita Corozal was not differentiated from any of the present-day Maya groups, and of ancient Maya groups it is only differentiated from Saki Tzul (p=0.020) (**Figs. S2, S3**).

Chi-squared results indicate an over-representation of haplogroups C and D in the most ancient groups. Haplogroups C5 and D4 were over-represented in Mayahak Cab Pek (standard residual C5 = 7.7957, p = 9.511*10^-13^; standard residual D4 = 10.163, p = 8.631*10^-22^), while C5, D1, and D4 were over-represented in Saki Tzul (C5 = 6.823, p = 9.927*10^-10^; D1 = 4.035, p = 0.004; D4 = 5.868, p = 3.941*10^-7^). Caracol, Santa Rita Corozal, and Mexico Colonial populations had haplogroup distributions more similar to the majority of the present-day Maya and comparative Native Mexican populations, with A haplogroups as the most common in these groups. Although A2 (the most frequent maternal haplogroup in the Americas today) is still the haplogroup of highest frequency in Santa Rita Corozal and in Caracol, these two ancient sites both exhibit over-representations of haplogroup A1 (Santa Rita Corozal = 6.202, p = 5.507*10^-8^, Caracol = 7.072, p = 1.938*10^-10^) (**Fig. S4**).

Networks show somewhat varying relationships between ancient and present-day Maya populations. The haplogroup C network shows one arm with individuals from the oldest sites: Saki Tzul and Mahak Cab Pek (**Fig. 4)**. Santa Rita Corozal is shown branching from a more basal inferred node (+1 more inferred node), most closely connected to the ancestral node (containing Yuto-nahua-and Oto-mague-speakers), the arm with the oldest samples, and one Tarahumara (Yuto-nahua language family) sample. The network for maternal haplogroup D also showed an arm exclusively containing samples from Mayahak Cab Pek and Saki Tzul. However, the samples from these oldest sites are shown to pair more closely to present-day Maya samples (Belizean Maya from Corozal in particular) than they are to the central node from which all the others radiate (**Fig. S6**).

Bayesian phylogenetic analysis recovered several well-supported internal tree nodes (posterior ≥ 0.8), particularly among deeper divergences, consistent with confident resolution of older branching events. However, a substantial proportion of nodes, particularly those representing more recent and population divergences, showed posterior support below this threshold, reflecting limited resolution for distinguishing recently diverged populations. Given this uncertainty in nodal support, we focus our interpretation on broadly supported topological patterns rather than specific divergence timing estimates, which we report for completeness in the supplemental materials.

Bayesian phylogenetic trees constructed from maternal haplogroup A show some ancient individuals were, interestingly, placed in clades that were not basal to the rest of the tree, while others ancient individuals were placed in basal clades. Ancient Caribbean individuals dating to the Ceramic Period (500 BCE – 1500 CE)^61,62^ cluster together with a couple of the Colonial Period individuals from Campeche and one from Mayahak Cab Pek (**Fig. 5**). This tree separated the ancient Maya individuals significantly, placing some individuals from Caracol basal to the rest of the tree, and one (Individual C53; Coverage 0.45x; **Table S2**) with individuals from Santa Rita Corozal in a clade with one present-day Totonaco individual from Veracruz, Mexico. Interestingly, the tree also split ancient individuals from Mayahak Cab Pek. Two individuals from Mayahak Cab Pek were placed in a clade exclusively composed of Maya speakers, while another was placed in a clade composed almost exclusively of Ceramic Period individuals from the Caribbean. The remaining samples from Mayahak Cab Pek, like the other ancient individuals, were scattered across clades (**Fig. 5**). Only one ancient individual was included in the haplogroup B tree (**Fig. S5**); this ancient individual from Santa Rita Corozal was placed basally to the modern samples. In the phylogenetic tree detailing mitochondrial relationships among maternal haplogroup C, ancient Maya individuals clustered together in a clade basal to all of the present-day individuals (**Fig. 4**). This clade included almost entirely individuals from Mayahak Cab Pek and Saki Tzul. This tree also showed one individual from Santa Rita Corozal, which was placed basal to all other clades (Individual SRC17; Coverage 1.02x; **Table S2**). Among haplogroup D, the Bayesian phylogenetic tree shows ancient individuals from Saki Tzul and Mayahak Cab Pek clustered together in a clade basal to the rest of the clades (**Fig. S6**).

### Maya in Northern Belize Show Evidence of Recent Maternal Gene Flow from Outside the American Double-Continent

We observe evidence of recent European, African, and Asian gene flow in the maternal lineages of present-day Maya people from the Northern Belizean district of Corozal, consistent with the colonial history of the region. Haplogrep results show five of the 110 individuals (4.5%) from the cohort of Belizean Maya from Corozal exhibited maternal haplogroups that were not autochthonous to the Americas. Two of these individuals exhibited L haplogroups (L0a1a2 and L1b1a), which are most common in sub-Saharan Africa today. One individual showed haplogroup H3w, which is most common in parts of Western/Northern Europe and Northwestern Africa. Two individuals exhibited maternal haplogroups (M30c and W3a1) found most commonly in Asia: M30c reaches its highest frequency in South Asia, while W3a1 is most commonly found in Northeastern Asia and the Middle East (**Fig. 6**).

**Figure 6.**
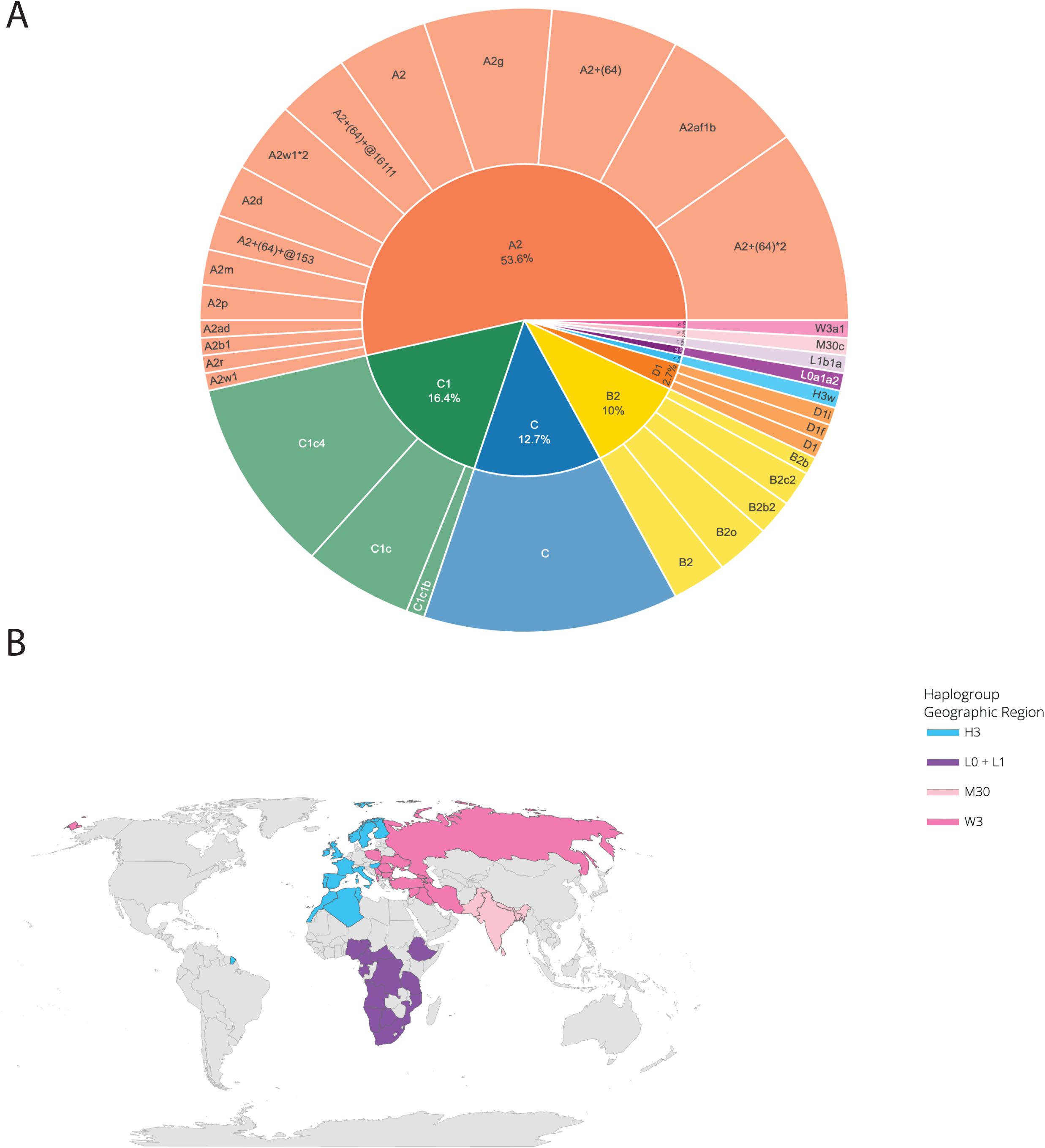
A) Haplogroup composition of present-day Belizean Maya study participants, B) Map showing geographic regions where non-Indigenous American haplogroups are common. Because specific location data is not always available for samples in the literature, modern country borders were used in this figure, noting that haplogroup distributions are not uniform throughout these countries.

## Discussion

### Maternal Lineage Sharing Across Linguistic Groups Reflects Deep Common Ancestry and Geographic Proximity Rather Than Language-Mediated Gene Flow

Despite linguistic variation between Mayan, Yuto-nahua, Oto-mangue, and Mixe-Zoque groups, maternal haplogroup diversity in this study does not cluster according to linguistic affiliation, a pattern consistent with recent whole mitogenome evidence from Mexican Indigenous populations more broadly^11^. The significant positive correlation between pairwise F_ST_ and geographic distance identified by Mantel test (r = 0.255, p = 0.005) suggests that geographic proximity is at least a contributing factor to the observed patterns of genetic similarity, though isolation by distance alone does not fully account for the patterns observed between geographically distant populations. When we considered mixed-language group clades containing at least one present-day Mayan speaker, Mayan speakers co-occurred in clades with Yuto-nahua speakers 81.5% (22/27) of the time, Oto-mangue speakers 70.4% of the time (19/27) and Mixe-zoque speakers 37.0% (10/27) of the time. This pattern of co-occurrence of contemporary Mayan, Yuto-nahua, and Oto-mangue speakers within shared mitochondrial subhaplogroup clades suggests that individuals from these linguistically distinct groups share common maternal ancestry, though the exact mechanisms of this shared ancestry cannot be determined from subhaplogroup membership alone.

Several demographic processes could produce this pattern, none of them mutually exclusive. First, these lineages may descend from a shared ancestral lineage that predates the divergence of the linguistic families themselves, in which case the genetic similarity would reflect deep common ancestry rather than recent contact or migration. This interpretation is consistent with the broad Mesoamerican distribution of haplogroups A, B, C, and D, which are thought to derive from the initial peopling of the Americas and would therefore be shared across many linguistically distinct descendant populations. Second, the pattern could reflect historical population movement and gene flow along known Mesoamerican corridors, particularly given previous research indicating that Maya centers regularly drew nonlocals from a variety of regions^17–22,63–66^.

Not all linguistic groups show equal levels of co-occurrence with Maya speakers, however. The relatively lower co-occurrence of Mixe-Zoque speakers compared to Yuto-Nahua and Oto-Mangue speakers may simply be reflective of the decreased number of Mixe-Zoque populations included in the study relative to Oto-Mangue and Yuto-Nahua populations. It is important to note that shared sub-haplogroup membership reflects ancestry at the level of the sub-haplogroup’s origin, which cannot necessarily be used to infer direct or recent gene flow between specific populations. As a result, patterns of gene flow between groups cannot be directly assessed without autosomal analysis. Taken together, the patterns of maternal lineage sharing documented here suggests deep shared maternal ancestry underlying linguistic diversity in Mesoamerica, shaped by the initial peopling of the continent and subsequent sustained demographic movement and gene flow rather than one where genetic differentiation aligns with linguistic differentiation^11^.

### Mitochondrial Haplogroup Distribution Suggests Broad Maternal Lineage Sharing Between Ancient and Present-day Maya Peoples with Increasing Genetic Differentiation Temporally

The pattern of low maternal lineage differentiation observed across the majority of Maya groups in this study is consistent with a broadly interconnected regional population that has maintained genetic continuity across much of the Holocene. Low F_ST_ values, non-significant Fisher’s exact tests, and central MDS placement across most Maya groups suggests that maternal haplogroup diversity in this region reflects a shared ancestral pool with high genetic connectivity rather than population-specific lineage histories. Our results are consistent with maternal lineage sharing between Maya groups extending since at least the Classic Period (250-∼900 CE)^16^, as evidenced by co-occurrence of ancient and contemporary Maya individuals in shared sub-haplogroup clades. However, this pattern may also be reflective of low mitochondrial diversity characteristic of Mesoamerican peoples more broadly, which derives from serial founder effects during the initial peopling of the American double-continent rather than from explicit gene flow between groups^67,68^. Increased sampling from time transects from the Archaic Period through the Pre-Classic Period and into the Classic Period will be necessary to determine continuity and lineage sharing across these time periods in more detail.

While the overall pattern across Maya groups is one of shared ancestry and low differentiation, closer examination of the temporal dimension reveals a story of gradual change. The progressive increase in F_ST_ between present-day Corozal and ancient sites as a function of mean site age is consistent with gradual population change following demographic replacement during the Archaic – Preclassic Periods. The over-representation of haplogroups C and D in the two oldest sites, which give way to the haplogroup A dominance characteristic of most present-day Maya and comparative Mexican Indigenous populations, suggests a shift in maternal lineage composition sometime between the Middle Archaic and the Classic Period, consistent with previous findings at Mayahak Cab Pek and Saki Tzul^60^. These previous findings indicate that the people occupying the Mayahak Cab Pek and Saki Tzul rock shelters ∼9.6-7.5 kya were genetically distinct from later individuals occupying the same rock shelters, who dated to about 5.5-3kya. The individuals dating to ∼9.6-7.5kya shared genetic ancestry with present-day South Americans and appeared to be more closely related to Anzick-1 than did the more recent group. In contrast, those dating to ∼5.5-3kya were skewed towards living Isthmo-Colombians on a PCA space and were more genetically similar to living Maya communities in Belize today. This is consistent with a South-to-North migration of Isthmian populations into the Maya area sometime between the occupation dates of these two groups^60^.

The evidence for human migration is broadly paralleled in genetic evidence from strains of maize, which was domesticated about 7kya in Southwestern Mexico. Maize agriculture quickly proliferated south into South America, differentiating in a variety of different strains as people selectively bred cultigens. Around 4kya, strains developed from South American maize lineages appear to move back into Belize, admixing with the strains that were already there^69^.Together, the genetic and archaeobotanical evidence points to the Northern Belizean Maya as part of a broader Maya population whose maternal lineage history reflects multiple waves of migration, gene flow, and in-place continuity stretching back to the initial peopling of the double-continent. This perspective provides essential context for interpreting the relationships between Corozal District and broader Mesoamerican linguistic and cultural groups, as discussed in the following section.

### Present-Day Belizean Maya from Corozal District Show Close Maternal Connections to Maya Groups in the Yucatán and with Contemporary Yuto-Nahua and Oto-Mangue Speakers

We highlight the Corozal Maya in this study not necessarily because they are a distinct cultural group, but because they have contributed novel samples in the study. Historically this is a Yucatec Mayan-speaking group with ancestral ties to the Yucatán via the Caste Wars of 1847 and before European colonialism. Our results indicate that present-day residents of the Corozal district who identify as Maya are a people that have maintained broad ancestral diversity without clear evidence of genetic drift or prolonged isolation, as evidenced by the absence of significant haplogroup over or under-representation in chi-squared analyses and low F_ST_ values relative to most present-day and recent ancient Maya populations. (**Fig. S1, S4**). The scattered placement of Corozal individuals throughout networks and trees, along with their central position in MDS analyses further suggests broad maternal lineage sharing with regional populations, with sustained connectivity between Northern Belize and broader lowland Maya populations across the Postclassic and into the present (**Fig. 2-5, Fig. S5-S6**).

Archaeological and historical context provide support for this interpretation, characterizing Northern Belize as part of a well-connected lowland Maya population rather than an isolated community. Northern Belize is known to have an extremely dynamic migration history, with major migration movements in direct response to political and demographic upheaval across the Maya world. The *Chilam Balam,* a group of documents written in the 17th and 18th centuries detailing traditional knowledge and histories of Maya groups in the Yucatan, tell of a major movement of people north from the Gulf Coast, Southern Lowlands, and Petén region (in what is now Northern Guatemala) into the Northern Lowlands^70,71^, now encompassed by the Mexican states of Campeche, Yucatán, and Quintana Roo (**Figure S8**); some of these same populations also moved back south at a later date^72^. These movements suggest that the genetic affinity observed between Corozal District and Yucatán Peninsula Maya populations may in part reflect these documented episodes of northward migration and resettlement, rather than just shared deep ancestry.

Many Maya polities experienced major depopulation during the Terminal Classic Period (750-∼900CE), with populations contracting around larger polities and their regional connections^34^. Although this period of depopulation occurred over the course of a few hundred years and was the culmination of a myriad of sociopolitical and environmental factors braided into an imbroglio still being parsed apart by archaeologists today, the results were a stark divergence from established political and ideological systems and depopulation of centers due to sociopolitical and economic instability, climatic stress, warfare, and displacement^14,73,74^. Comparisons of residential structures during this time period suggests a demographic decline of an estimated 75-95% across the Maya Lowlands, with considerable regional variation^12,50,75^. However, many Northern coastal cities, including those in Northern Belize, grew in population and maintained connections throughout the region, thriving off of extensive maritime trade routes well into the Postclassic Period^17,76–80^.

Northern Belize thus represents one important locus of the social, political, and economic reorganizations that characterized many Maya peoples during the Postclassic, facilitating continued demographic exchange with groups across the lowland Maya region and potentially beyond. The broad maternal lineage connectivity that characterizes present-day residents of the Corozal District has deep roots in the pre-Colonial demographic history of Northern Belize.

### Possible Maternal Genetic Continuity Between Late Postclassic Santa Rita Corozal and Present-Day Maya in Corozal District, Despite Demographic Continuity

Among the ancient Maya sites included in this study, Santa Rita Corozal is of particular interest given its geographic proximity to present-day Maya communities living in Corozal District and the site’s well-documented role as a regional capital during the Postclassic Period^17,76,77,81^. The genetic relationship between Santa Rita Corozal and present-day Maya in the Corozal District therefore offers a direct opportunity to assess maternal lineage continuity within a single geographic locality across several hundred years.

The extremely low F_ST_ value between present-day Maya individuals in Corozal and ancient Maya individuals at Santa Rita Corozal is consistent with matrilineal continuity within the Corozal District across the Late Postclassic. Based on the data thus far, we find no evidence of a major matrilineal discontinuity between the Late Postclassic group at Santa Rita Corozal and the present-day Maya community in the Corozal district, a finding consistent with other noted instances of genetic continuity in other parts of the Maya region, namely that ancient residents of Chichen Itza show a high genetic affinity with modern residents of nearby Tixcacaltuyub^51^.

This finding is also consistent with archaeological understanding of the area. Santa Rita Corozal was a Northern Belizean Maya city which was consistently occupied since the Middle Preclassic Period, but which reached its peak occupation in the Postclassic Period^82,83^. The city has long been a bridge between the Northern and Southern Maya Lowlands and maintained a uniquely consistent occupation and proportion of emigrants throughout these time periods^17,23,78,82,84^. Santa Rita Corozal is located very close to Corozal Town, though the genetic continuity between the two sites has never been formally tested or quantified.

Together, the observed low degrees of genetic differentiation and archaeological evidence of demographic occupation through the Late Postclassic and into the present is consistent with the persistence of some maternal lineages in the area despite the political and social disruptions of the Colonial Period, including the shift from Spanish to British colonial rule, the Caste War, and the fairly recent struggle for Belizean independence from the British Crown, achieved in 1981. These results indicate that genetic continuity and demographic connectivity may not be mutually exclusive processes. While consistent with other documented instances of Maya genetic persistence, our results should be interpreted with caution given the limitations of mitochondrial data and small sample size among the Santa Rita Corozal ancient samples. To fully characterize the continuity between these two sites in the future, mitogenomes from additional time-transects at Santa Rita Corozal should be included, as well as autosomal loci.

### Caracol’s Maternal Genetic Distinctiveness May Reflect Limitations in Sampling and Preservation Rather Than Broader Population Structure

Caracol presents a more complex interpretive picture than Santa Rita Corozal, in part because the genetic signal observed at this site is difficult to reconcile with its political and economic prominence during the Classic Period. Unlike the pattern of low differentiation at Santa Rita Corozal, Caracol shows significant F_ST_ differentiation from nearly all other ancient and present-day Maya populations, a finding that requires consideration of both the archaeological context of the samples and the limitations of the dataset. One of the largest Maya cities at its peak, with extensive causeway networks and political dominance over neighboring polities including Tikal^85,86^, Caracol might be expected to show relatively low differentiation from surrounding populations due to its high connectivity. The fact that our limited data indicates it does not suggests that political connectivity may not always translate into detectable maternal gene flow, a topic which merits further investigation in the future.

A critical consideration here is that our interpretation is based on seven samples, three of which exhibit less than 1x mitogenome coverage (**Table S2**). While there are enough diagnostic SNPs to make haplogroup calls for these samples (**Table S2**), the Caracol results therefore cannot be taken as representative of the city’s demographic history as a whole and warrant further investigation with larger sample sizes and further spatial coverage within the site. In addition, the very limited number of individuals from Caracol included in this study derive from residential mortuary contexts in the mixed-status residential groups (nicknamed Jack, Chak, and Tuna) near urban administrative node known as the Puchituk Terminus, 3 km distant from the city’s epicenter^87–90^. As a result, these individuals could have represented migrants from surrounding rural areas or middle-status inhabitants who may have had less access to the long-distance networks centered at the political core^91^. If the total population of Caracol was more demographically diverse due to long-distance political and economic connections, the very limited samples included in this study may capture a more locally contained subset which is not representative of the broader urban population, explaining the higher differentiation observed despite the city’s political reach. This interpretation would be consistent with growing evidence from other Mesoamerican urban centers such as Teotihuacán for significant intra-site demographic heterogeneity between elite and non-elite residential contexts, though Caracol presents an urban and administrative structure distinct from Teotihuacán^44,92,93^. Thus, explicit hypothesis testing would be required to understand the relationship between city structuring and genetic heterogeneity at Caracol during these time periods.

Despite these limitations, the Caracol results raise interesting questions about the relationship within and between distinct sectors of Classic Period cities. If peripheral residential populations at Classic Period urban centers were found to be more locally derived than epicenter populations^94^, this would have important ramifications for how we interpret aDNA data from urban Maya contexts more broadly and underscores the importance of sampling strategy in genomic studies of ancient cities.

### Present-Day Maya Individuals from Corozal Exhibit Evidence of Recent Admixture with African, European, and South Asian Populations

Among the present-day Belizean individuals sampled from the Corozal District, a small proportion of maternal lineages belong to haplogroups that originate from regions outside of the Americas. Specifically, we observe the presence of haplogroups L (African in origin), H and W (European and Central Eurasian in origin), and M30 (South Asian in origin) in our Corozal cohort^95–101^. While these lineages represent a minority of the total mitogenomes in this study, their presence is consistent with the well-documented history of the Colonial Period in Northern Belize and reflects the broader pattern of sex-biased gene flow documented across Mesoamerican populations more generally^3^.

Spanish colonialism began in the region in 1527 CE, when conquistador Francisco de Montejo was given official authority by the Spanish crown to take control of the Yucatán peninsula and the island of Cozumel; only the northern part of the Yucatan Peninsula was brought under control by 1542 CE^102^. These events, coupled with the earlier spread of European-introduced disease (especially smallpox) across the region, initiated a second, expedited, period of catastrophic demographic decline in the Maya region. This demographic catastrophe, taking place within about 50 years, resulted from the combined effects of epidemic disease, forced labor systems, warfare, and Spanish policies of forced resettlement (*congregaciones*) that disrupted Indigenous lifeways^103^. Maya communities resisted Spanish control vehemently, fleeing these areas of forced resettlement, with the Itza successfully maintaining freedom from Spanish rule until 1697 CE^39,104,105^. Some Maya towns in Northern Belize like Lamanai grew during this period, as people fled from areas of conflict^39,106^, reinforcing the role of Northern Belize as a refuge and demographic receiving area during periods of regional upheaval.

British colonialism in Belize followed a different trajectory, focused primarily on timber extraction and relying on enslaved African labor rather than Indigenous exploitation, though Maya communities in the region nevertheless experienced population decline and displacement^40^. During the late 1630s, a group of pirates known as Baymen settled on the coast to start logging operations. Over time, the Baymen intensified their efforts harvesting logwood and later mahogany, and increasingly relied on bringing in enslaved Africans from nearby British colonies to work in the logging operations^107^. There is also a documented history of South Asian indentured labor in Belize during the period following the abolition of slavery in 1838. Following emancipation, the British colonial government sought to replace the sugar plantation labor force with indentured workers primarily from the Indian subcontinent^108^. This colonial labor system reshaped population movements and demography in the region and provides important historical context for the African and South Asian maternal lineages that were identified in this study.

Northern Belize was again transformed demographically in 1847, serving as a major haven for Maya refugees escaping the Caste Wars in the Yucatán^52,104,109^. An estimated 7,000 migrants crossed from Mexico to Northern Belize in the first year of the Caste Wars alone^109^. Between 1848 and the formal end of the conflict in 1901, the influx of refugees fundamentally altered the demographic and economic structure of Northern Belize, transforming the region from a timber extraction economy into one based on sugar cane cultivation and further diversifying the population of Corozal District^109,110^.

Taken together, the maternal lineages found in living people in Corozal who identify as Maya present a complex but uneven mosaic of Indigenous, European, African, and South Asian genetic ancestries reflective of the region’s history of Indigenous continuity, colonial violence, forced and voluntary migrations, and local Maya resilience. The observed pattern of maternal lineage sharing is broadly consistent with documented patterns across Mesoamerica, where the Colonial Period introduced African, European, and in some cases Asian maternal lineages into Indigenous populations through the mechanisms of the Transatlantic Slave Trade, forced labor systems, and indentured migration^3,111^. However, the specific composition of gene flow in the present-day Corozal Maya originating from outside of the Americas reflects the unique colonial structure that characterized Belize relative to the broader Mesoamerican region. Under Spanish colonial rule, which governed the majority of Mesoamerica including the Yucatán, Guatemala, and Mexico, admixture was characterized primarily by the introduction of European paternal lineages through the Spanish settler population and African maternal lineages through the Transatlantic Slave Trade. This pattern produced the heavily sex-biased admixture signal documented across Mexican and Guatemalan Maya populations, in which European ancestry is predominantly transmitted through the Y chromosome while Indigenous American and African ancestry is predominantly transmitted through mtDNA^3^. Present-day Guatemalan Maya STR-based analyses show near complete Indigenous ancestry despite substantial European Y-chromosome ancestry^112^, reflecting the highly sex-biased nature of Spanish colonial admixture in that region, while Yucatán Maya show higher levels of European admixture overall, consistent with more intensive post-Columbian gene flow in that region compared to Highland Guatemala^42^.

Northern Belize, by contrast, was heavily shaped by British colonial structures in addition to those imposed by the Spanish, which focused on timber and sugar extraction using enslaved African and later South Asian indentured labor rather than Spanish settler colonialism^104,108^. As a result, the maternal lineages introduced into the Corozal District from outside of the Americas reflect this distinctly British colonial demographic footprint, rather than the European maternal lineages more commonly introduced through Spanish settler colonialism elsewhere in Mesoamerica. Critically, the overall proportion of maternal lineages originating from outside of the Americas in the Corozal Maya cohort remains small, consistent with the broader pattern across Mesoamerica of Maya maternal lineage resilience in the face of colonial disruption^3,112^. This pattern reflects both the sex-biased nature of colonial admixture and the demographic persistence of Indigenous communities despite catastrophic population decline.

### Further Analysis of Autosomal Variation Among Maya Population May Elucidate Recent Gene Flow Between These Groups

Overall, we observe a substantial amount of shared maternal ancestry across groups included in this study. Such lineage sharing may be indicative of sustained gene flow between these groups. Although some patterning by linguistic family is evident, we observe no clear genetic structuring among the mitochondrial genomes of Maya peoples included in this study. Similarly, we observed no clustering of Maya groups separating them genetically from other Mexican Indigenous populations. The placement of the present-day Belizean population from Corozal that identify as Maya across a wide variety of phylogenetic positions relative to both ancient and contemporary mitogenomes, rather than in self-contained clusters, suggests either ongoing or historical gene flow with neighboring populations, and/or retention of ancestral diversity predating cultural, political, and linguistic differentiation in the region. The limited resolution of mitochondrial data for recent population divergences highlights the need for future genome-wide autosomal analyses of these populations, which will add resolution to our understanding of regional connections between Maya people throughout history and provide greater power to resolve fine-scale population structure and more recent demographic events, along with the possible sex-bias in these patterns.

## Materials and Methods

### Ethical Approach

This project adheres to all existing legal and ethical standards in the United States and Belize for genomic and paleogenomic research. This study was conducted with approval and support of the Institute of Archaeology in Belize, and approval by George Mason University’s IRB (No. 1919667-1). This research was co-developed and conducted by members of the research team, including local Maya peoples and the Maya-led NGO To’one Masehualoon, which is based in Corozal Town and focuses on the preservation of Yucatec Maya culture, language, and knowledge of the region. Research objectives in this study were developed jointly with Maya collaborators in Corozal. Laboratory and data governance protocols were also co-developed with the express purpose of ensuring cultural appropriateness and respect for Maya Ancestors and their remains. Project co-directors and US-based members of the research team have visited the participant communities in Belize at least once annually since 2022 to return results to study participants and to the local community. Results of the contemporary mitochondrial and autosomal DNA analysis presented here were returned to communities in 2025.

### DNA Extraction, Library Preparation, Sequencing, and Quality Screening

Skeletal and dental samples were collected from 16 Ancient Maya Ancestors from the ancient cities of Caracol (N=7) and Santa Rita Corozal (N=9) (**Table S1, S2**). Ancient DNA was extracted in a dedicated paleogenomic cleanroom following a modified version of Dabney et al. (2013)^113^ and using methods for respectful treatment of Maya Ancestors developed in collaboration with our Maya community partners. After aDNA extraction, dual-barcoded and dual-indexed double stranded Illumina libraries were prepared and screened using a modified version of Rohland et al. (2015)^114^. Resulting libraries from all samples were enriched using the myBaits® Expert Mito probes (Arbor Biosciences) and standard protocols targeting the complete mitogenome. Libraries from ancient samples were sequenced on an Illumina HiSeq 2000. Resulting ancient sequence data were cleaned, mapped, and variants called using the EAGER Pipeline^115^. Coverage and contamination estimates for the libraries are shown in **Table S2**.

Saliva samples were collected from 110 contemporary residents of the Corozal district of Northern Belize who identify as Maya using Oragene OGR600 kits from DNA Genotek (**Table S1**). DNA was extracted from saliva samples and screened for quality using standard protocols with Genotek prep-IT-L2P reagents. Dual-indexed Illumina libraries for contemporary DNA extracts were prepared via the Enzymatic Fragmentation Kit 2.0 from Twist Bioscience using a modified protocol. Contemporary libraries were pooled and enriched using the Twist Bioscience mitochondrial panel targeting the whole mitogenome. Enriched libraries were sequenced on an Illumina NovaSeq X. Sequence data was prepared for analysis following GATK Mitochondrial Single Nucleotide Variant (SNV + Indels) best practices^116^.

Newly sequenced mitogenomes were combined with previously published ancient mitogenomes from the Allen Ancient DNA Resource (AADR)^59^ and present-day mitogenomes from Metabolic Analysis of an Indigenous Sample (MAIS) biobank^54–56^ (**Table S1**). The AADR was screened for whole mitogenome samples in the Maya region (Yucatan region of Mexico, Belize, El Salvador, Honduras) with at least 3x coverage and were maternal haplogroups native to the region (A, B, C, and D). This yielded samples from the colonial town of Campeche in Mexico^117^ (N=7) and Archaic Period (8000 – 2000 BCE)^16^ samples from the Belizean rock shelters Mayahak Cab Pek^118^ (N=15) and Saki Tzul^118^ (N=5). To add comparative ancient samples, we included from the AADR Ceramic Period (500 BCE – 1500 CE)^61,62^ samples from the Caribbean with the maternal haplogroup A2+64, a maternal haplogroup that is also common for Mixe-Zoque speakers and Maya linguistic families^4^. This yielded 16 additional samples dating to the Ceramic Period from the Dominican Republic^61^ (N=11) and the island of Curacao^61^ (N=1) (**Table S1**). These samples were all target-captured on the 1240K SNP array.

The MAIS biobank, initially collected to investigate metabolic disease in Native Mexican (NM) populations, was included in this study as a comparative dataset^2^ (**Table S1**). The biobank includes whole mitogenome data for 72 populations across Mexico, spanning 10 language families, and totaling 598 individuals^2^ (**Table S1**). These whole mitogenome samples were amplified via long range PCR and sequenced via the Ion PGM TM System^2^.

The final dataset utilized in this study included a total of 724 individuals spanning Mexico, Belize, and the Caribbean across the last ∼9,000 years (**Table S1**).

### Population Genetic Analysis

Maternal haplogroups were determined using Haplogrep 3^119^. Arlequin 3.5.2.2^120^ was used to calculate population molecular pairwise F_ST_ values, and Fisher’s exact tests. Significance was determined with 10,000 permutations. Population pairwise F_ST_ matrices were constructed to assess genetic similarity between each pair of populations. Fisher’s exact tests assessed if maternal haplogroup frequencies differed between each pair of populations. To characterize haplotype diversity and assess the genealogical relationships among mitochondrial lineages, we constructed median joining networks using PopArt v1.7^121^, with epsilon set to 0 to produce the most parsimonious network.

To test for over- or under-representation of maternal haplogroups in each population relative to the total dataset, chi squared tests of independence were performed using the base R stats package on observed counts for each population/haplogroup combination. Standardized Pearson residuals were tested against a standard normal distribution of residuals with FDR correction via the “p.adjust” function to see if any of the haplogroups are over- or under-represented in each population, where the sign of the standard residual is indicative of over (positive) vs under (negative) representation.

Multidimensional Scaling (MDS) was utilized to visualize population pairwise sequence difference in three-dimensional space. The pairwise F_ST_ matrix was converted into a distance matrix using the MDS function in “smacof” R package^122^. Both metric and nonmetric (ordinal) solutions were evaluated; the ordinal solution was chosen based on lower stress, indicating a better fit.

A Mantel test was performed using the R packages “vegan” and “geosphere” to assess potential association between geographic (Haversine) distance and genetic distance^123,124^. Pairwise F_ST_ values were used as genetic distances and geographic coordinates were derived from published latitude-longitude information where available. Significance was assessed using a one-tailed Pearson correlation with 9999 permutations. Coordinates for populations from samples generated in this study were the estimated centroid of the site from which the samples were taken, and in the case of the Belizean individuals identifying as Maya from Corozal, the coordinate we used was the centroid of the Corozal district. This allowed us to treat study participants who resided in 10 different towns around the district as one population. Additionally, and more importantly, this helped to ensure the privacy of the participants. The Nahuatl Hidalgo (NAH_HGO) population from the MAIS cohort (n=7) was excluded from the Mantel test as coordinates were not available in the original source data^125^ or associated supplementary materials. Any populations with only one sample in the population were also excluded.

To test evolutionary relationships among mitochondrial lineages, Bayesian phylogenetic trees were built for maternal haplogroups A, B, C, and D, using the two individuals from Corozal with L0 and L1 haplogroups as an outgroup to root the trees. Trees were constructed and annotated using BEAST suite, with XML files prepared using BEAUti, Bayesian analysis conducted in BEAST, trees annotated with TreeAnnotator and then visualized in R using “ggtree” and “treeio” packages. Bayesian analysis used whole mitochondrial genomes with the site model GTR+G, as determined by PartitionFinder to best fit our dataset. Additional parameter settings were consistent with Fu et al. (2013) (whole mitogenome with no partitions; ambiguities box checked; strict clock; substitution rate of 2.67x10^-8^; coalescent constant population prior)^126^. Markov Chain Monte Carlo (MCMC) settings were set for a chain length of 100 million, storing every 1000 iterations. Burn in was set to 10 million, with 10 initialization attempts. Tip dates were assigned for each sample in years BP (common ancestor heights), using dates reported in the AADR when available. If dates were reported in relative terms (from ceramic dating) we used the mean date of that range. ESS values were visualized in Tracer v1.7.2 to ensure all were above 200.

When visualizing trees, we identified haplogroup clades containing present-day Maya speakers to assess language group composition of the nodes within those clades. Language family classification followed that previously recorded for the MAIS dataset, with the understanding that some study participants are descended from speakers of that language and may not speak that language themselves. Additionally, pre-Maya individuals from Mayahak Cab Pek and Saki Tzul were included in the Ancient Maya language family due to their location in what is now referred to as the Maya region. Clades were defined by identifying the most recent common ancestor (MRCA) of all samples sharing a given sub-haplogroup designation as assigned by HaploGrep3. For each haplogroup tree, the MRCA node of each sub-haplogroup represented in that tree was identified using the “getMRCA()” function from the “ape” package in R, and all tips descending from that node were treated as members of that clade^127^. Only clades in which the descendent tips represented at least two distinct linguistic families were retained for annotation and composition analysis. Boxes identifying those clades on phylogenetic trees were colored by the dominant language family within the clade, and stacked bar plots were constructed to visualize the relative composition of language families within the clade. Clades were named based on their sub-haplogroup designation

## Data Availability

Whole mitogenome data generated in this study are available to researchers upon request pending review by the co-directors of the Corozal Population History Project (RWAS, AWR, AJR, LCS) and the members of the board of To’one Masehualoon.

## Acknowledgements

We would like to thank all of the participants and Maya communities for participating in this study. In particular, we would like to thank the Maya-led NGO T’one Masehualoon for their continued partnership in the development and ongoing implementation of this work. This project was funded in part through a

Level 1 grant (EC-93180R-22) from the National Geographic Society to RWAS and a grant from the National Science Foundation (2444410) to AWR and RWAS. The archaeological samples utilized in this study were collected from Santa Rita Corozal and Caracol through archaeological projects funded through a variety of sources that included NSF and the Alphawood Foundation to AFC and DZC. The funders had no role in the planning or execution of this study, nor in the drafting of this manuscript.

