## Supplemental Materials for "Ancient Mitogenomes from Belize Provide Insights into the Mobility and Migration of Ancient Communities in Mesoamerica and the Caribbean"

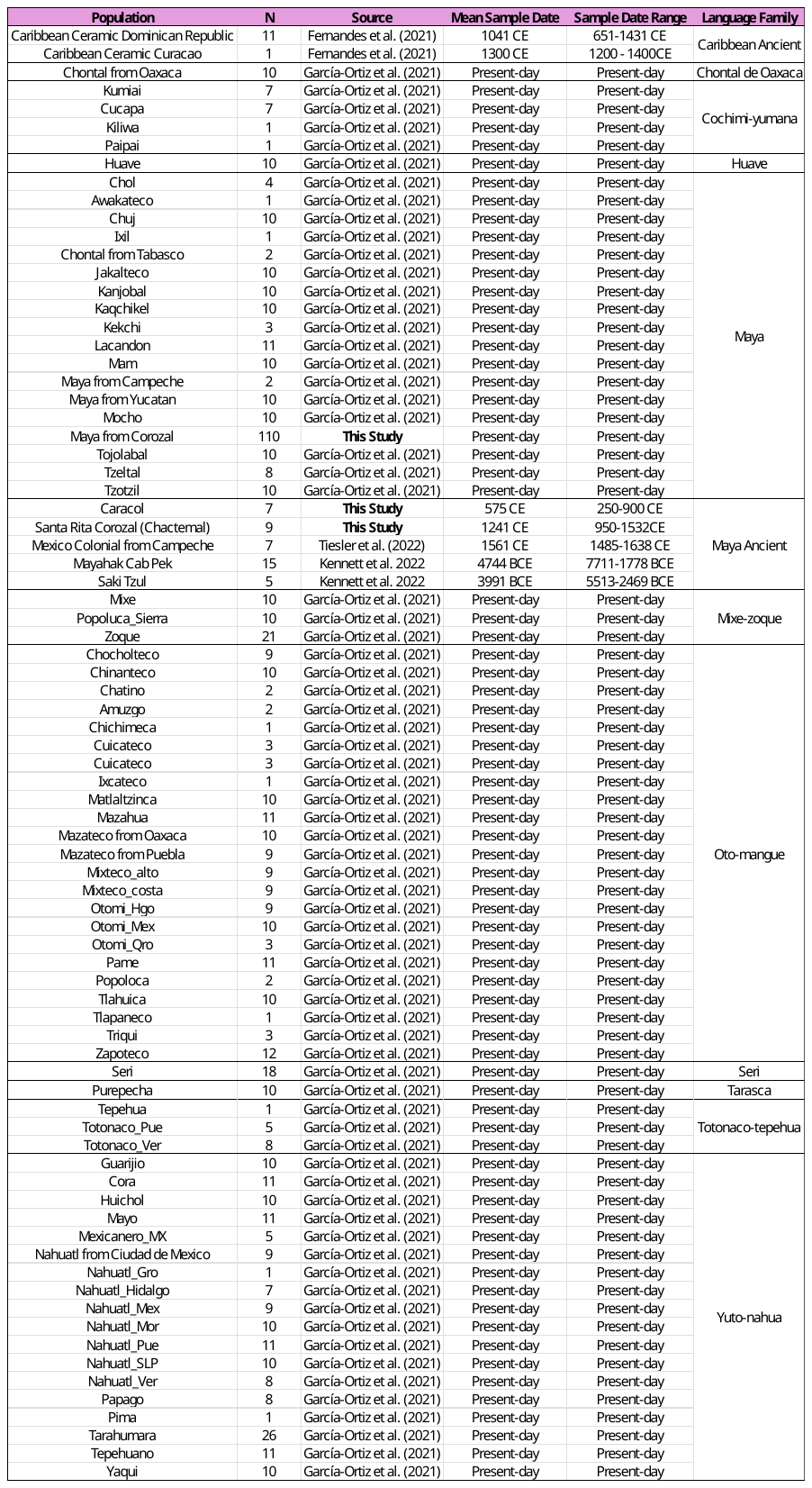
**SUPPLEMENTAL MATERIALS**

**Table S1**. Dataset

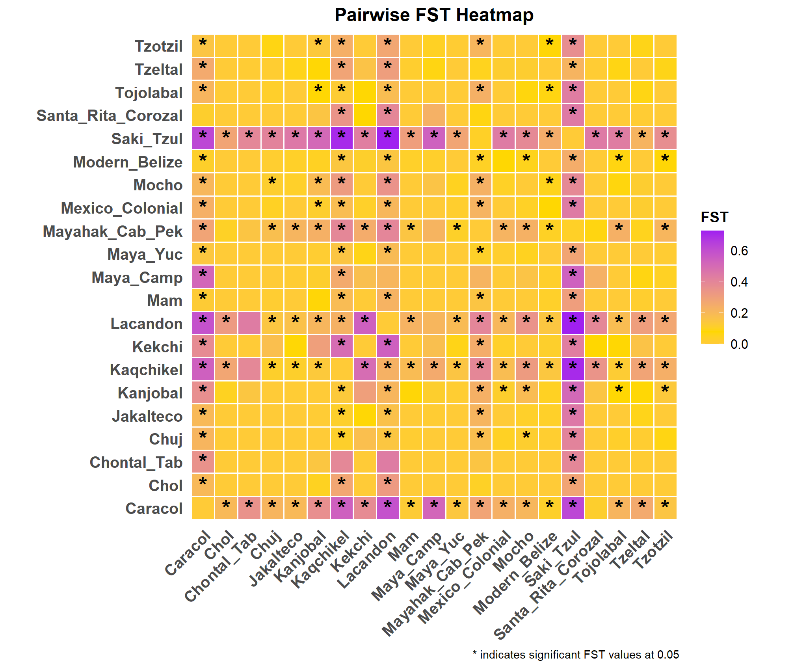

**Figure S1.** FST heatmap showing pairwise relationships between populations in the Maya language family. * Denotes significant FST values at alpha = 0.05.

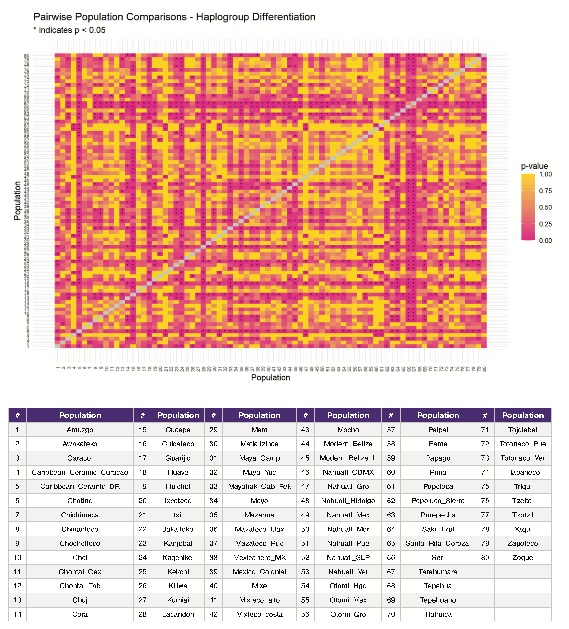

**Figure S2**. Fisher's exact tests of differentiation between all pairs of populations included in this study. * Denotes significant differentiation at alpha = 0.05.

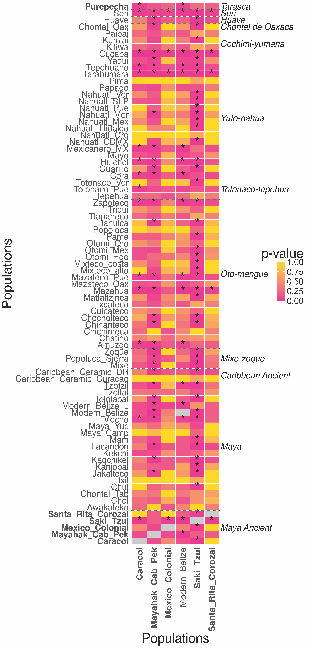

**Figure S3**. Fisher's exact tests of differentiation heatmap showing pairwise relationships between populations. This heatmap is reduced to emphasize relationships within and between ancient Maya and present-day Belizean Maya from Corozal populations.

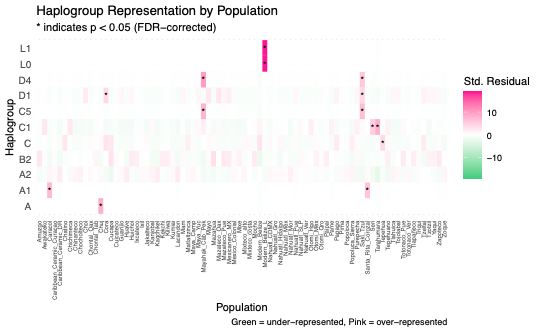

**Figure S4**. Chi-squared tests showing haplogroup over- or under-representation of haplogroups in each population. * Denotes significance at alpha = 0.05.

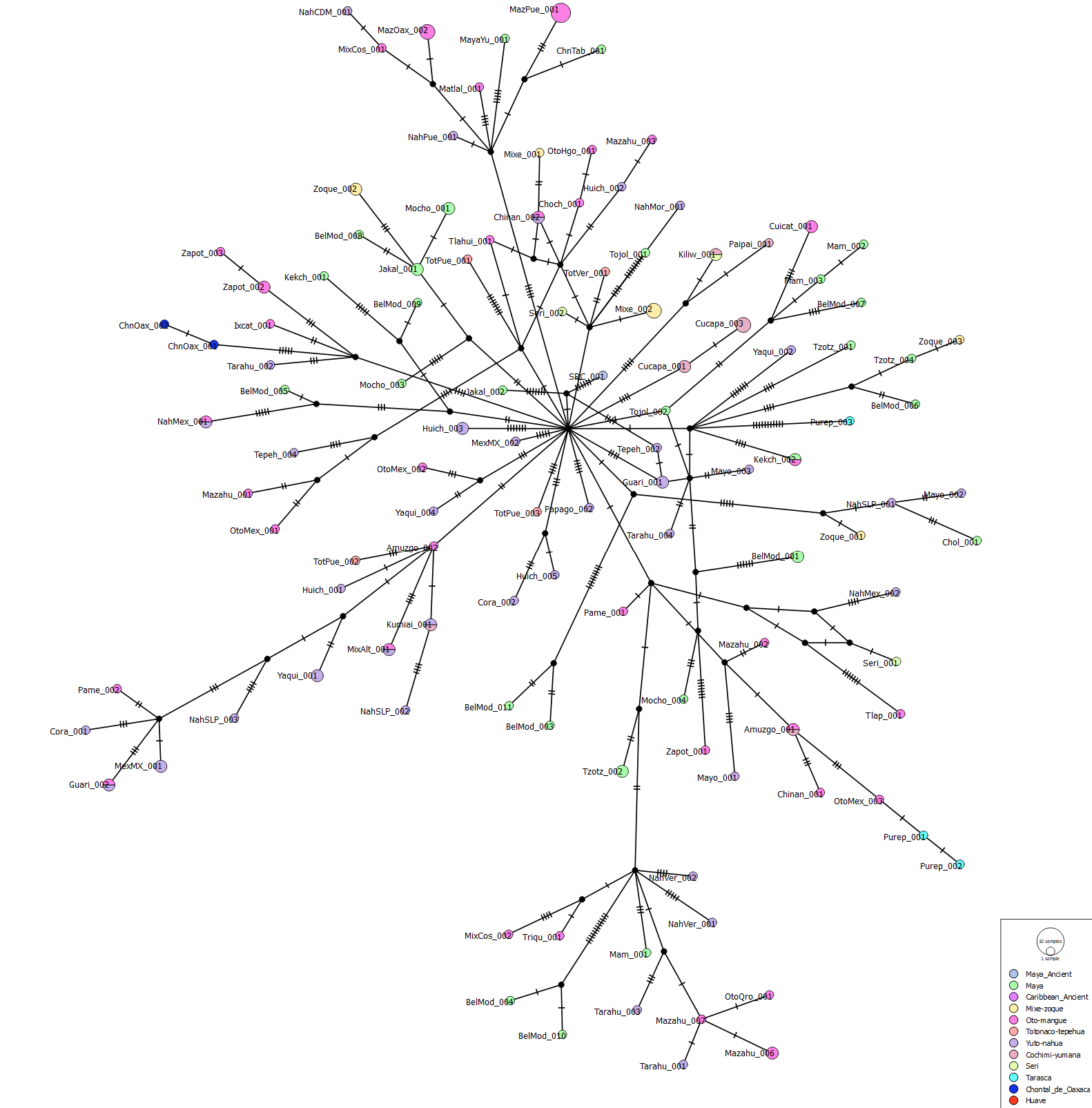

**Figure S5**. Haplogroup B median joining network

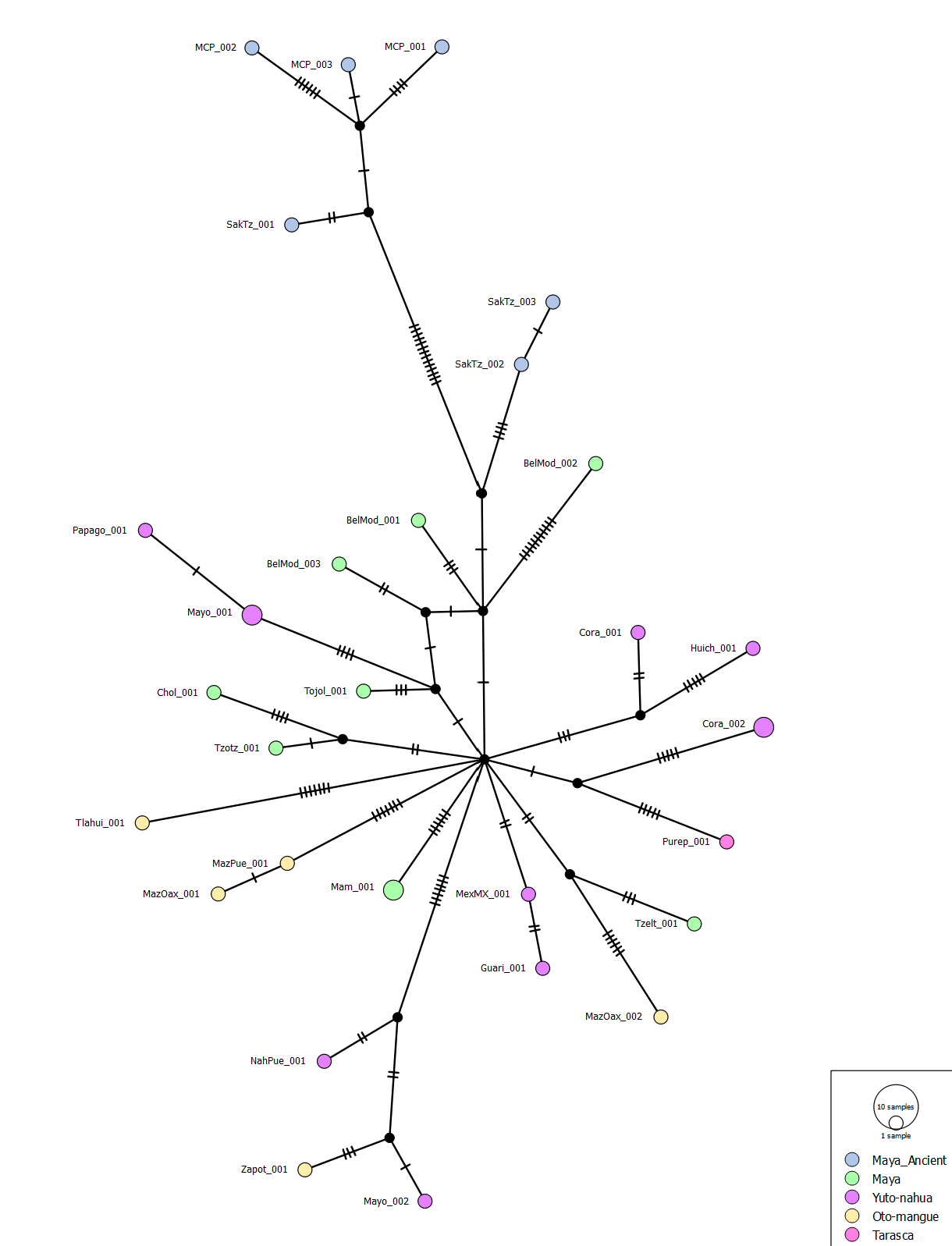

**Figure S6**. Haplogroup D median joining network.

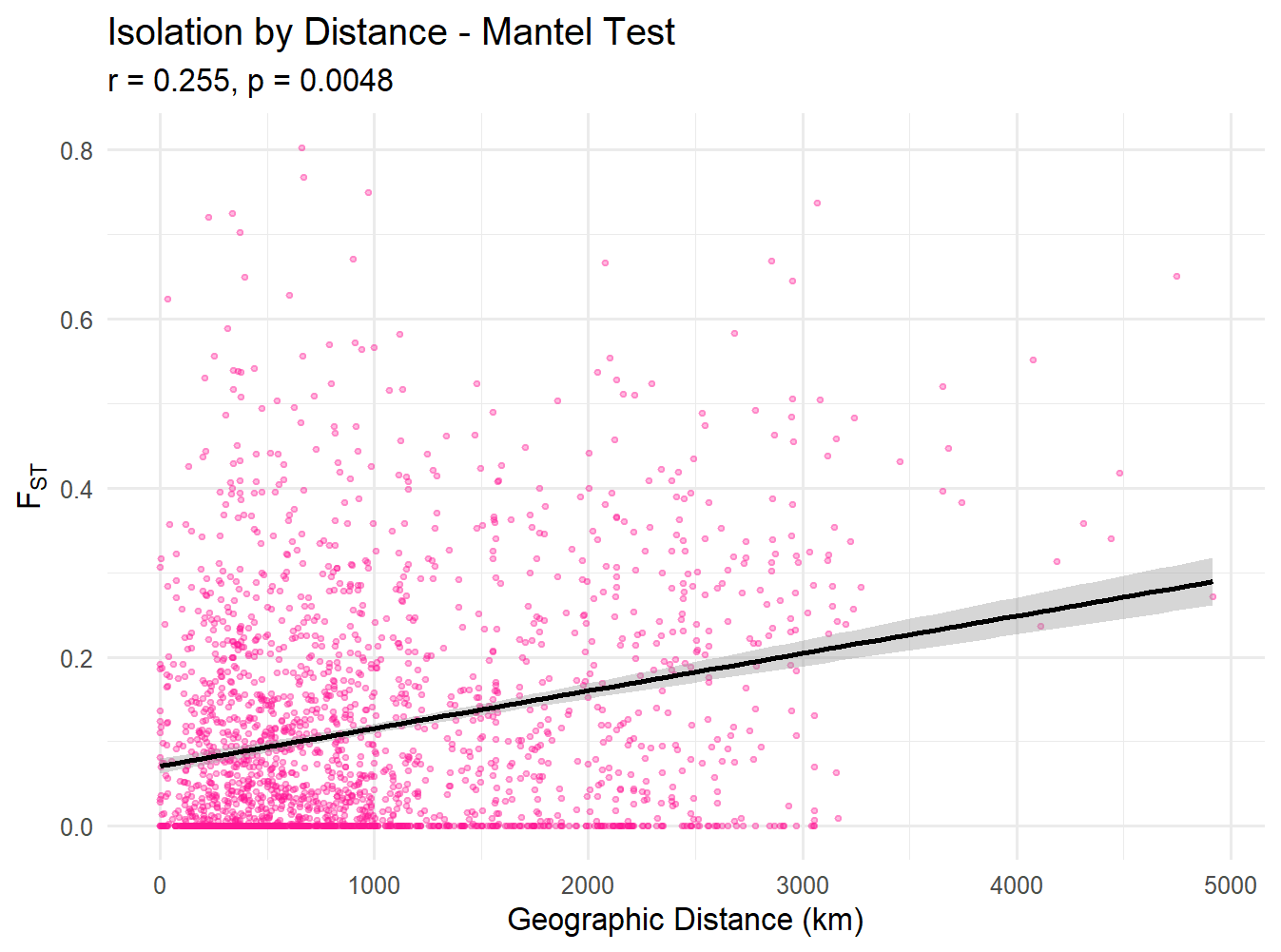

**Figure S7**. Mantel test results

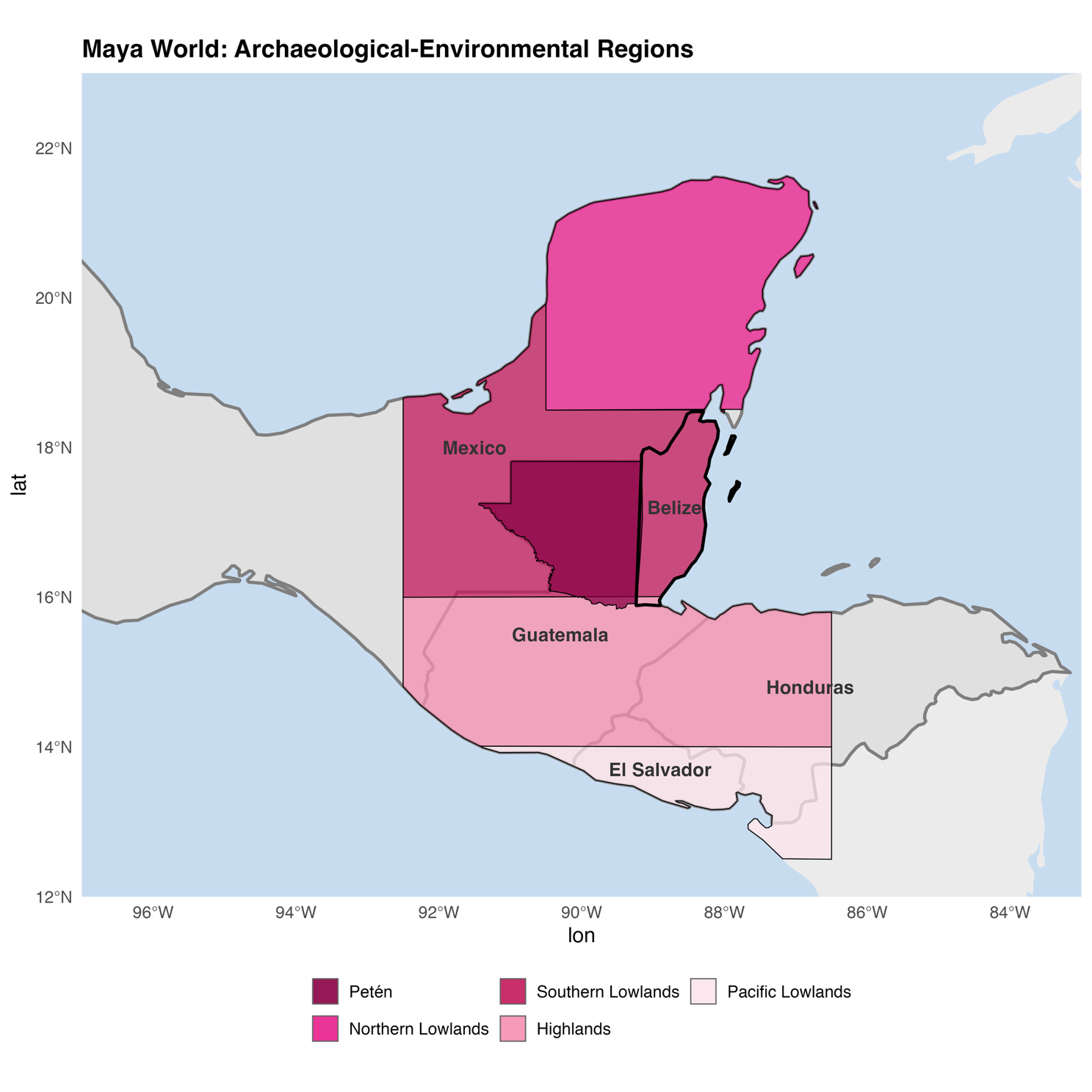

**Figure S8**. Map of archaeological-environmental zones in the Maya region.

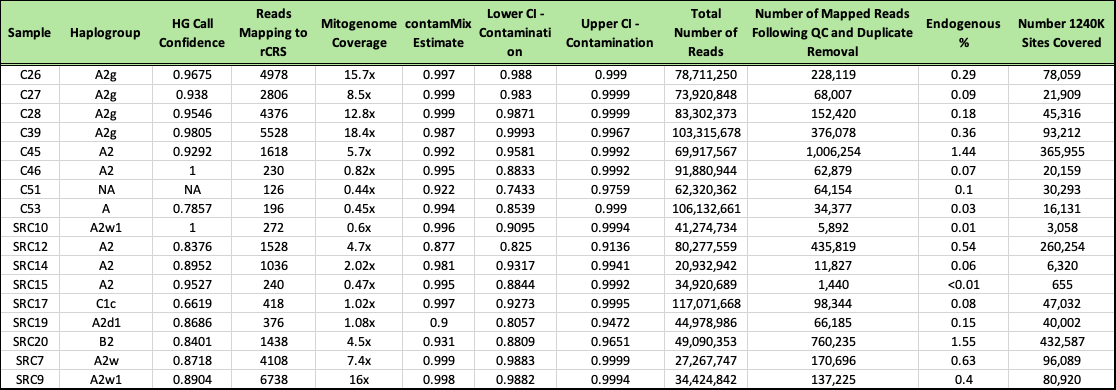

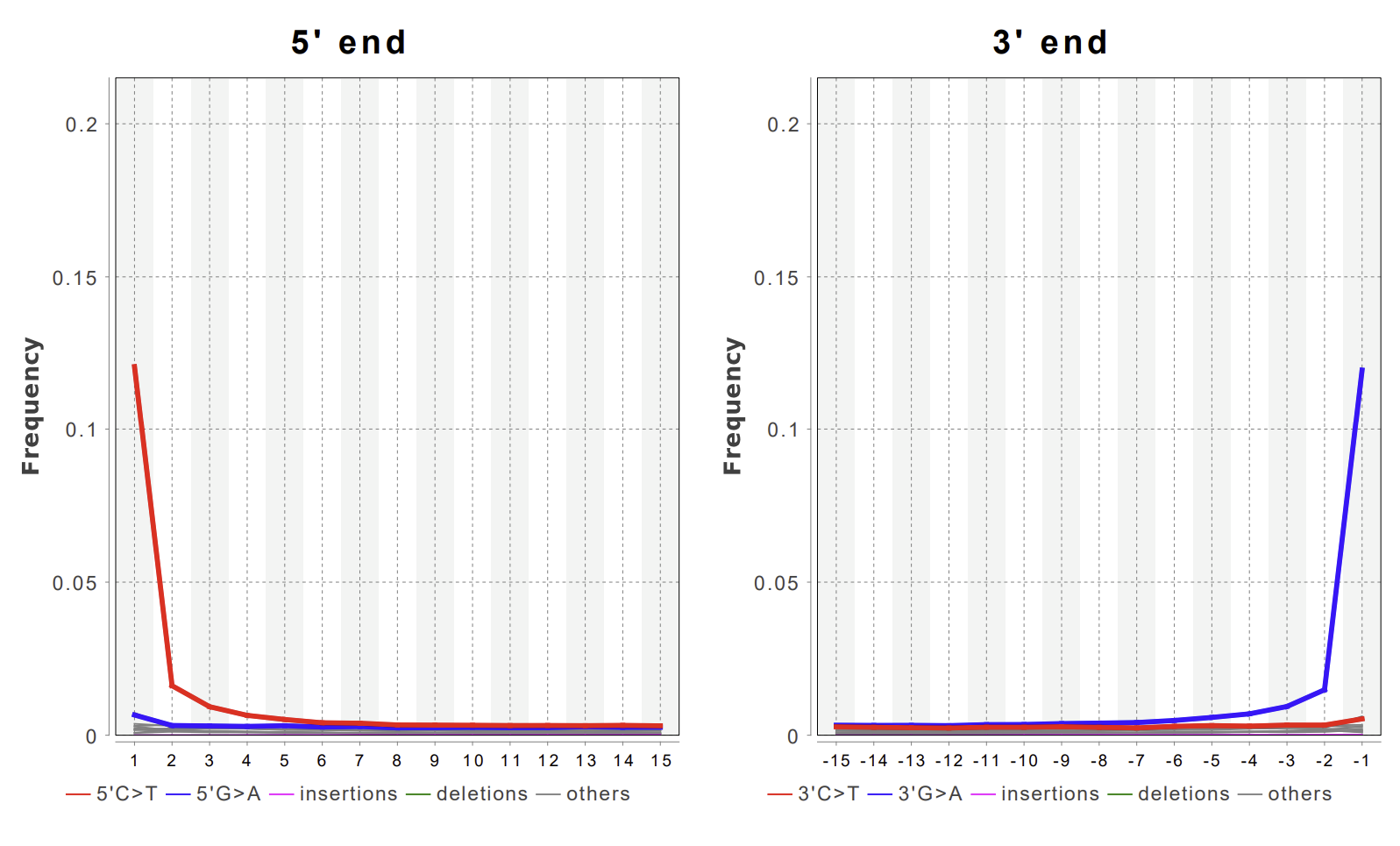

**Table S2.** Quality information for DNA extraction, library construction, sequencing results, and haplogroup calling for ancient samples generated by this study. Sample codes beginning with "C" denote samples from Caracol, and those beginning with "SRC" are from Chactemal (Santa Rita Corozal).

**Figure S9**. MapDamage plot from individual C45 (Caracol)
